# Mass cytometry data integration methods reveal rural-urban gradient of immune profiles across geography

**DOI:** 10.1101/2025.09.05.673697

**Authors:** Koen A. Stam, Jeremia J. Pyuza, Mikhael Manurung, Dicky L. Tahapary, Taniawati Supali, Marloes M.A.R. van Dorst, Ahmed Mahfouz, Mohammed Charrout, Simon P. Jochems, Wesley Huisman, Marion König, Yvonne C.M. Kruize, Marouba Cisse, Ibrahima Diallo, Moustapha Mbow, Maria Yazdanbakhsh, Wouter A.A. de Steenhuijsen Piters

## Abstract

The human immune system strongly varies across populations and is shaped by a wide range of host and environmental factors. As such, a rural compared to urban lifestyle has previously been associated with baseline differences in immune profiles, which can impact outcome of vaccination. The question whether there is a immune signature that associates with rural living irrespective of ancestry or geography has been addressed here along with which analytical methods are most suitable. Using three mass cytometry datasets, we studied shared and population-specific immune characteristics of healthy rural- or urban-living Indonesian, Senegalese and Tanzanian adults and urban-living Europeans. After harmonized preprocessing and quality control, 75.4 million cells were integrated using four different data-integration methods. Among these, CytoNorm with *in silico* references performed best, revealing shared differences in the differentiation state within the lymphocyte compartment that distinguishes rural from urban-living individuals. Differentiated CD4^+^ T cells expressing CTLA-4, PD-1 and ICOS characterized rural living, as did CD4^+^ T cells with high CD161 expression in combination with CRTH2 and GATA3. In the innate compartment, CD56⁻ CD16⁺ NK cells with low CD38 and CD11c expression were expanded in rural living subjects. Together, our results demonstrate that, using a harmonized analytical pipeline and machine learning-based integration, large-scale mass cytometry data can be combined across diverse cohorts to identify shared immunological hallmarks of rural living. These signatures may provide targets for modulating immune responses in future vaccine studies.

## Introduction

The invention of vaccines for the prevention of infectious diseases has been one of the greatest scientific innovations. Each year, millions of deaths worldwide are prevented due to childhood immunization (Li et al., 2021). Although the use of vaccines has been a remarkable medical breakthrough, research findings from across the globe have shown striking variation in vaccine performance across different populations. Lower responses have been observed in populations living in low- and middle-income countries (LMICs) compared to high-income countries (HICs). Reduced vaccine immunogenicity, often referred to as *vaccine hypo-responsiveness*, has been described across multiple vaccine platforms (van Dorst et al., 2024), including vaccines against BCG tuberculosis, ebolavirus and yellow fever (YF) (Muyanja et al., 2014; Natukunda et al., 2024). The lower immunogenicity can translate into lower vaccine efficacy, which has been observed for among others, rotavirus and malaria vaccines. For example, attenuated malaria vaccine protects 92% of vaccinated participants in Europe (Epstein et al., 2017), but drops to 27% in Africa (Sissoko et al., 2017), Pre-vaccination immune signature profiles play a critical role in shaping vaccine responsiveness (Fourati et al., 2022; Hagan et al., 2022; HIPC-CHI Signatures Project Team & HIPC-I Consortium, 2017; Kotliarov et al., 2020; Tsang et al., 2014). For example, a study revealed a shared pro-inflammatory transcriptional signature linked to higher serum antibody responses one month post-vaccination across 13 vaccines (Fourati et al., 2022). Of note, these integrative studies are mainly based on blood transcriptional profiles of healthy individuals living in HICs, lacking data on populations in LMICs. Even within LMICs, baseline immune activation and as such vaccine-induced immune responses vary (Temba et al., 2021). A recent study on Tanzanian adults showed that urban living was related to a stronger pro-inflammatory profile compared to individuals adopting a rural lifestyle (Temba et al., 2021).

Baseline immune state is shaped by a range of factors, including age, heritable (genetics) and non-heritable factors (e.g. vaccination, nutrition (Temba et al., 2021), CMV or helminth infections, epigenetics and microbiota composition) (Brodin & Davis, 2017; Brodin et al., 2015) In particular, non-heritable and lifestyle factors co-occur with the degree of urbanization, possibly explaining differences in baseline immune signatures and responsiveness to vaccines. Immune profiles related to helminth-infected, rural-living Indonesians were characterized by expanded frequencies of T helper 2 and regulatory T cells expressing CTLA-4, a marker associated with the suppressive function of these cells. Upon deworming treatment, CTLA-4-expressing regulatory T cells in rural-living Indonesians decreased, but not reaching the low levels of uninfected European individuals. (de Ruiter et al., 2020) As baseline immune states are likely driven by a multitude of underlying factors, part of which may not even be known at this point, it is therefore important to map the immune networks of populations living along the rural-urban gradient in LMIC settings. Doing so, we may be able to divert unfavorable environmental exposures on baseline immune states by targeting the downstream immune processes. This approach would be particularly successful if immune hallmarks are shared between low-responding populations, despite the fact that environmental drivers leading to these immunological characteristics may be different.

To characterize immune profiles along the rural-urban gradient in detail, we integrated mass cytometry datasets across three studies, using different analytical methods to assess their performance. Immune profiles from 172 healthy adults from urban and rural sites (Indonesia, Tanzania and Senegal) and from an urban area in The Netherlands, were used allowing us to both assess shared and population-specific immune profiles across geography.

## Results

### Study participants

For this study, we included 172 participants from three studies (Senegal, CapTan and SugarSpin) conducted across four countries (Senegal, Tanzania, Indonesia and the Netherlands; Figure 1A, Table S1). All participants were healthy adults. The median age was 26 years (range, 17 – 55 years) and 52% of participants was female (Table S2). For non-European countries, samples were collected from individuals living in urban or rural areas, which previously was shown to be associated with distinct cellular immune profiles (de Ruiter et al., 2020; Manurung et al., 2025; Pyuza et al., 2024). All participants had always lived in their respective residence areas. Helminth infection status was tested and summarized, showing different prevalences in the rural areas (Table 1).

**Figure 1.**
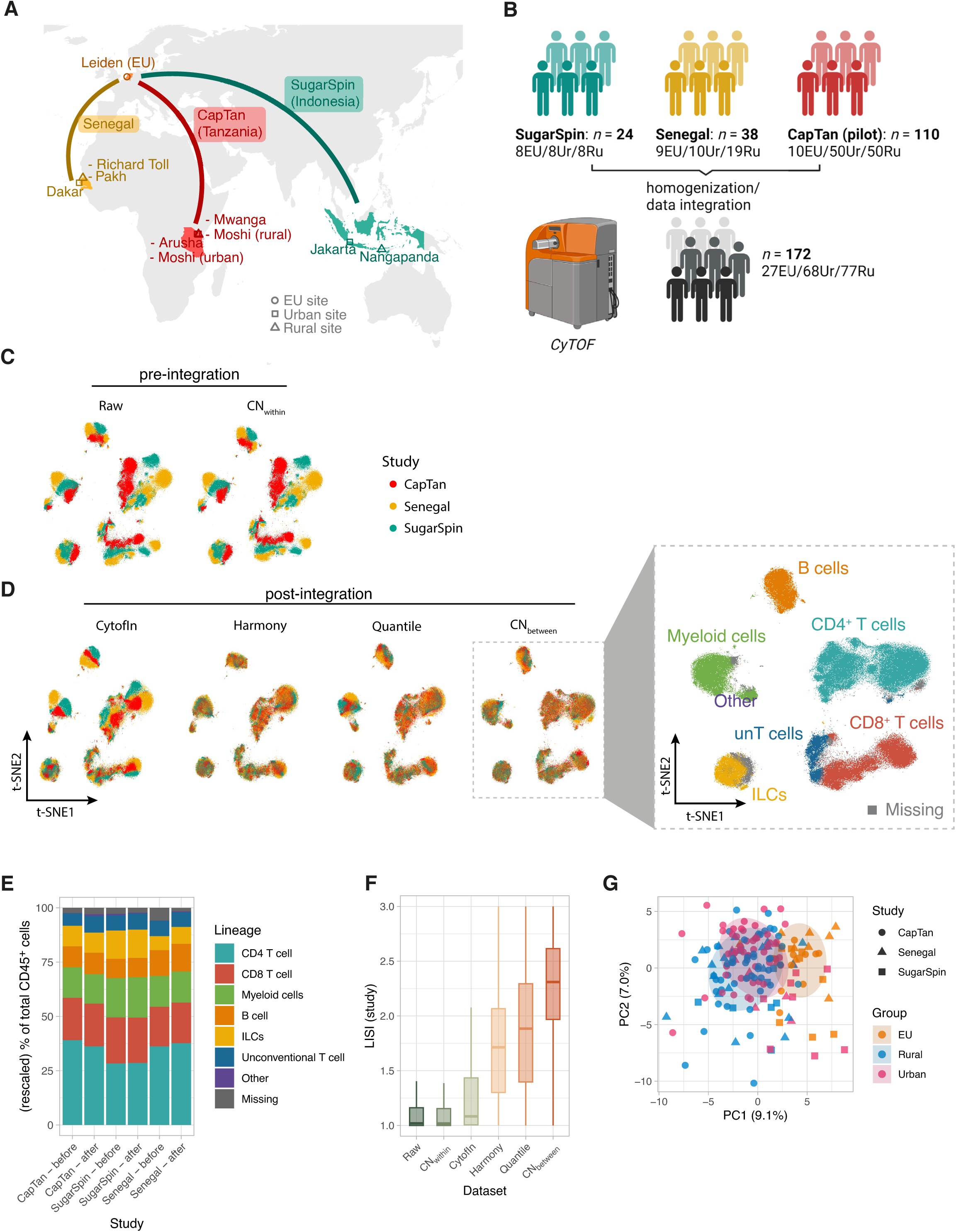
CytoNorm successfully integrates three mass cytometry datasets. A) Geographical map of studies and study sites. Symbols indicate EU, urban or rural sites. B) Graphical representation of sample numbers and the study design. C-D) t-distributed Stochastic Neighbor Embedding (t-SNE) visualizations for each dataset C) pre- [‘raw’ and CN_within_] and D) post-integration [CytofIn, Harmony, quantile and CN_between_-normalisation]; *n* = 50k random cells/study); cells are colored according to study (panels) and lineage (zoomed-in area). E) Stacked bar chart of lineage frequencies pre- (CN_within_; ‘before’) and post-integration (CN_between_; ‘after’) for each of the studies. Each bar section represents the mean frequency of that cell lineage for the corresponding group. F) Local inverse Simpson index (LISI) scores of study across datasets (pre- and post-integration). G) Principal component analysis (PCA) was applied to cell cluster frequencies (centered and scaled to unit variance) of individual samples (*n* = 172). Each participant is reflected as one data point. Data points are colored based on group (European, EU; Rural and Urban). The shape of datapoints indicates the study (CapTan, Tanzania; Senegal and SugarSpin, Indonesia studies). Ellipses reflect the data spread at a level of confidence of 68% (one standard deviation in a normal distribution).

### Generating an integrated mass cytometry dataset accounting across studies

From all individuals, PBMC samples were available, from which we profiled a total of 183 million immune cells using mass cytometry and largely identical panels (Figure 1B, Table S3). Using these data, we generated an integrated dataset of mass cytometry studies conducted across geographical locations. First, raw (bead-normalized) data were preprocessed through a unified pipeline (Figure S1). The resulting dataset (75.4 million cells) was used without further preprocessing (‘raw’) or were subjected to within-study batch-correction using CytoNorm (CN_within_) or between-study batch-correction (CytofIn (Lo et al., 2022), Harmony (Korsunsky et al., 2019), quantile or CytoNorm[CN_between_] (Van Gassen et al., 2020)). Global differences in the performance of integration methods were visualized using t-distributed Stochastic Neighbor Embedding (t-SNE). These t-SNE-plots clearly underscored the need for data-integration methods, revealing strong separation according to cell lineage and study in non-integrated and within-study batch-adjusted datasets. In contrast, batch effects (largely) disappeared in the integrated datasets (CytofIn, Harmony, quantile and CN_between_), and cells mostly separated based on cell lineage (Figure 1D-E).

To quantify integration performance in more detail, we calculated the local inverse Simpson index (LISI), which is a diversity score that measures how well studies are mixed in the local neighborhood of each cell (Korsunsky et al., 2019). LISI showed marked differences between integration methods, with the highest median LISI-scores in CN_between_-adjusted data (median 2.39, IQR 2.08-2.67; Figure 1F), indicating superior integration compared to other methods. Similarly, *k*-nearest-neighbor batch-effect test (kBET)-scores indicated best integration was reached for the CN_between_-adjusted data (Figure S2).

### Automated annotation of cell clusters and comparisons before and after integration

Given the superior integration metrics, we decided to proceed with the CN_between_-adjusted data for downstream analyses. Data were clustered using self-organizing maps (SOM; 15 × 15 grid) and meta-clustering was performed for a range of *k-*values (total number of clusters). Based on cluster-level inverse Simpson indices (ISI) across European samples, we determined that the optimal number of clusters was k = 125, balancing a peak in the ISI score with a high number of clusters to capture granular differences in the immune system (first local peak in median ISI across clusters with k ≥ 100; Figures S3 and S4). As a first exploratory analysis, we performed principal component analysis (PCA) on the cluster frequencies. The first two principal components explained 16.1% of the variance in the data (Figure 1G). Strongest separation was observed along the first principal component, where Europeans clearly clustered apart from non-Europeans, which was verified using PERMANOVA-tests (*R^2^*= 8.9%, *p*-value = 0.0001, permutations restricted within study strata). Despite this, a residual effect of study ID was visible, mostly along the second principal component (PERMANOVA *R^2^* = 5.4%, *p*-value = 0.0001). We next assessed overall cell cluster diversity, both assessing the number of clusters present at cell frequency > 0.1% and the evenness of the distribution of these clusters. We found that rural-living individuals overall have a more diverse immune profile compared to both urban-living non-Europeans and Europeans (Shannon index; linear model, adjusting for study; *q*-value 0.01 and 1.6 × 10^−10^, respectively; Figure S5A), which was particularly apparent in the CD4^+^ T cell compartment (Figure S5B). This difference may be explained by higher variation in environmental exposures related to rural lifestyle, resulting in a more heterogeneous immune cell signature in rural living individuals.

Next, cell clusters were annotated using an automated and standardized marker-based algorithm. In brief, the algorithm derives for each cluster whether a given marker is positive, negative or neutral, based on the intensity distribution across all clusters for that marker. For each cluster a label is assigned based on a set of predefined ‘rules’ (Table S4). To assess the effect of integration on cell type annotations, we employed the automatic annotation scheme to both the unintegrated and integrated data. We assessed the effect of the integration and the ability to preserve biology by comparing cell type annotation of the integrated data to the unintegrated data using the F1-measure (Figure 2A-C). The effects of integration with CN_between_ were minimal at the lineage level, with the median F1-measure (measure for accuracy of cell labels post-integration compared to pre-integration labels) across lineages ranging between 0.96-0.98. However, at the subset-level, F1-measures dropped, with median subset-level scores of 0.77, 0.78 and 0.82 for the CapTan, Senegal and SugarSpin study, respectively (Figure 2C). We found that lower-scoring subsets, such as CD4^+^ CM T cells (F1-scores 0.66-0.73), are split across CD4^+^ CM, EM and naive cells after integration (Figure 2A). Furthermore, although different CD8 T cell subsets (i.e. CD8 EM, CM and EMRA) could be discerned before integration in each study, a portion of these cells were annotated as ‘CD8 T cell’ after integration (especially in Captan and Senegal, F1-score 0.34 and 0.35, respectively). ‘CD8 T cell’ is a general label given to cells within the CD8 T cell lineage that lack clear negative or positive markers to further distinguish between specific subsets (Table S4). On the other hand, specific cell phenotypes, including B cells, classical monocytes, ILC2s, and pDCs, showed consistently high F1 scores (>0.8) and thus good integration across all studies (Figure 2C). Of note, subsets with lower F1 scores were often defined by markers tagged with metal isotopes that differed between studies (e.g. for CapTan, Senegal and SugarSpin studies, Gd_155_, Pt_198_ or Cd_113_-isotopes were conjugated to CD45RA-antibodies, respectively; Table S3). This confirms that differences in panel design between mass cytometry datasets poses a challenge upon data integration.

**Figure 2.**
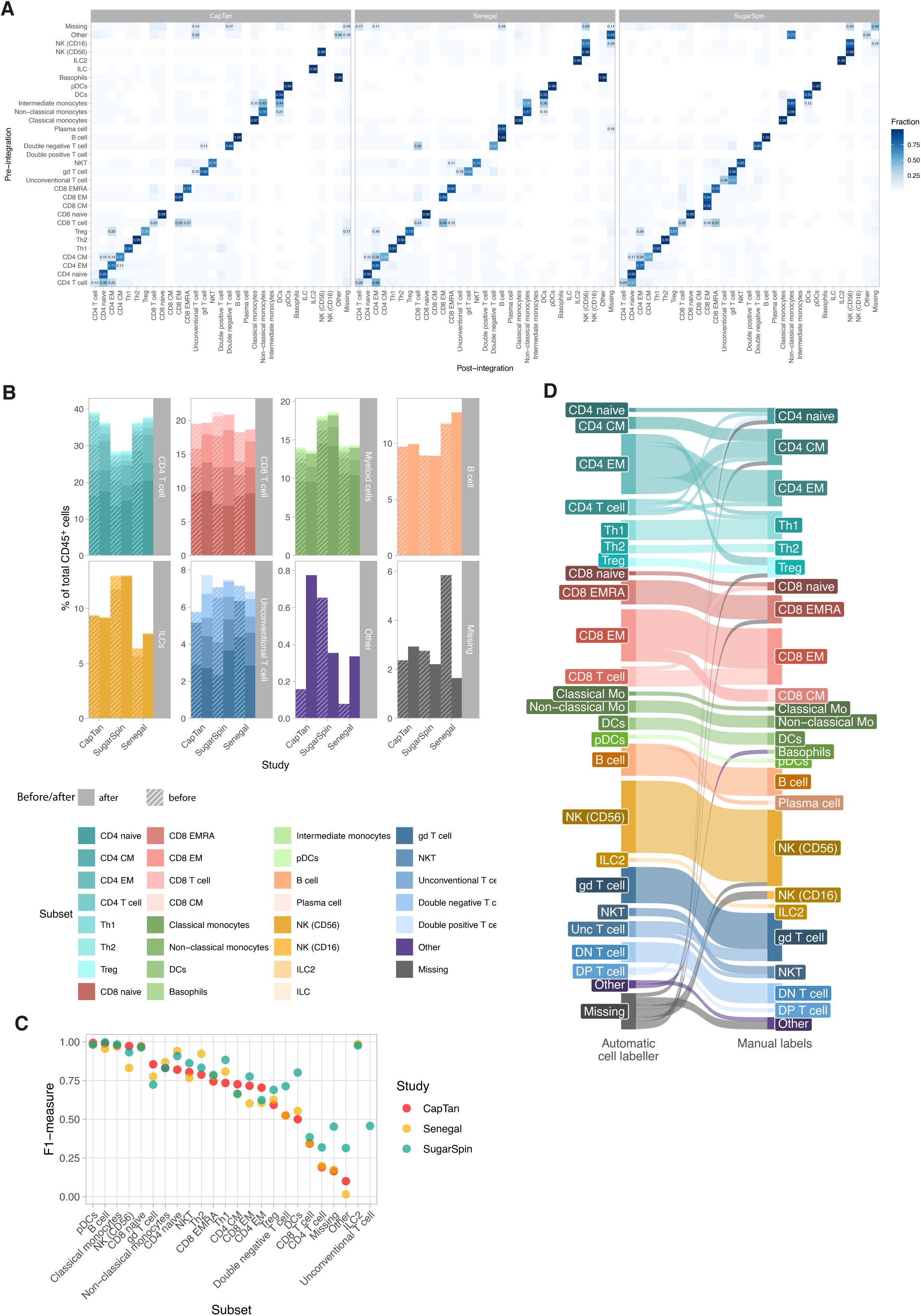
Automated post-integration subset-level annotations of cellular immune profiles adequately capture cellular immune profiles. A) Confusion matrices for each study, showing the subset-level annotations generated by the automatic labelling algorithm before and after data integration. Colours and values in the cells reflect how a given pre-integration label is distributed across post-integration labels (‘fraction’). B) Stacked bar plot showing cell frequencies (number of cells/total CD45^+^ cells) for each subset. Subset labels were based on automatic cell labelling before (CN_within_) and after (CN_between_) data integration. Cell frequencies before [striped] and after [filled] integration are shown and facetted according to cell lineage. C) F1 scores (harmonic mean of the precision and recall) as a measure of labelling accuracy was computed for each cell subset (comparing data before [truth] and after [prediction] integration). D) Alluvial diagram showing the results of which subsets were re-labeled after the automatic cell labeler.

Next, to evaluate cell populations in the context of urban–rural differences, we manually reviewed the annotations generated by the automatic cell labeler to ensure accurate and consistent phenotyping. (Figure 2D, Table S5). Of the 125 clusters, 90 retained their original labels after manual review, meaning no changes were made to their assigned subset or lineage. Among the remaining 35 clusters, the labeler was unable to annotate 9. Of these, 2 were relabeled as “other,” indicating that their phenotype could not be matched to a meaningful lineage or subset (e.g., negative for all markers).

Consequently, only 7 of the 125 clusters required manual labeling from scratch. For the other 24 clusters, manual edits involved minor refinements to better specify the phenotype. For example, clusters initially labeled with general categories such as “CD4 T cell” or “CD8 T cell” were updated to more specific phenotypes, such as “CD8 Naive” or “CD8 EM” (Figure 2D).

### The immune composition of rural living individuals is emphasized by CD4^+^ T cell differentiation

To study variation in the immune cell composition between populations, a generalized linear mixed model was used to assess how integrated immune cell populations differed between urban and rural-living non-European individuals (Figure 3, Figure S6-S7). As a secondary analysis, we compared immune profiles of these individuals to urban-living Europeans, hypothesizing this group would represent the most urban signature, allowing us to study rural-urban differences across a collective gradient. This gradient was classified based on the significance and direction of differences between 1) rural and urban-living non-Europeans (Ru - Ur), 2) rural-living individual non-Europeans and Europeans (Ru - EU) and 3) urban-living non-Europeans and Europeans (Ur - EU). Significant differences in all comparisons (1., 2. and 3.) were referred to as a ‘full urban > rural gradient’ or ‘full urban < rural gradient’ depending on the direction of the effects (Figure 3A-B). Overall, we found 11 clusters following the full rural > urban gradient (whereby cluster frequencies were highest in rural and lowest in Europeans). In contrast, none of the clusters exhibited a full urban < rural gradient (Figure 3A-B). Also considering partial urban – rural gradients (comparisons 1. and 2. or 2. and 3. significant) and limited urban – rural differences (comparison 1. significant), we identified in total 20 clusters and 3 subsets. (Figure 3D; Figure S6-S7).

**Figure 3.**
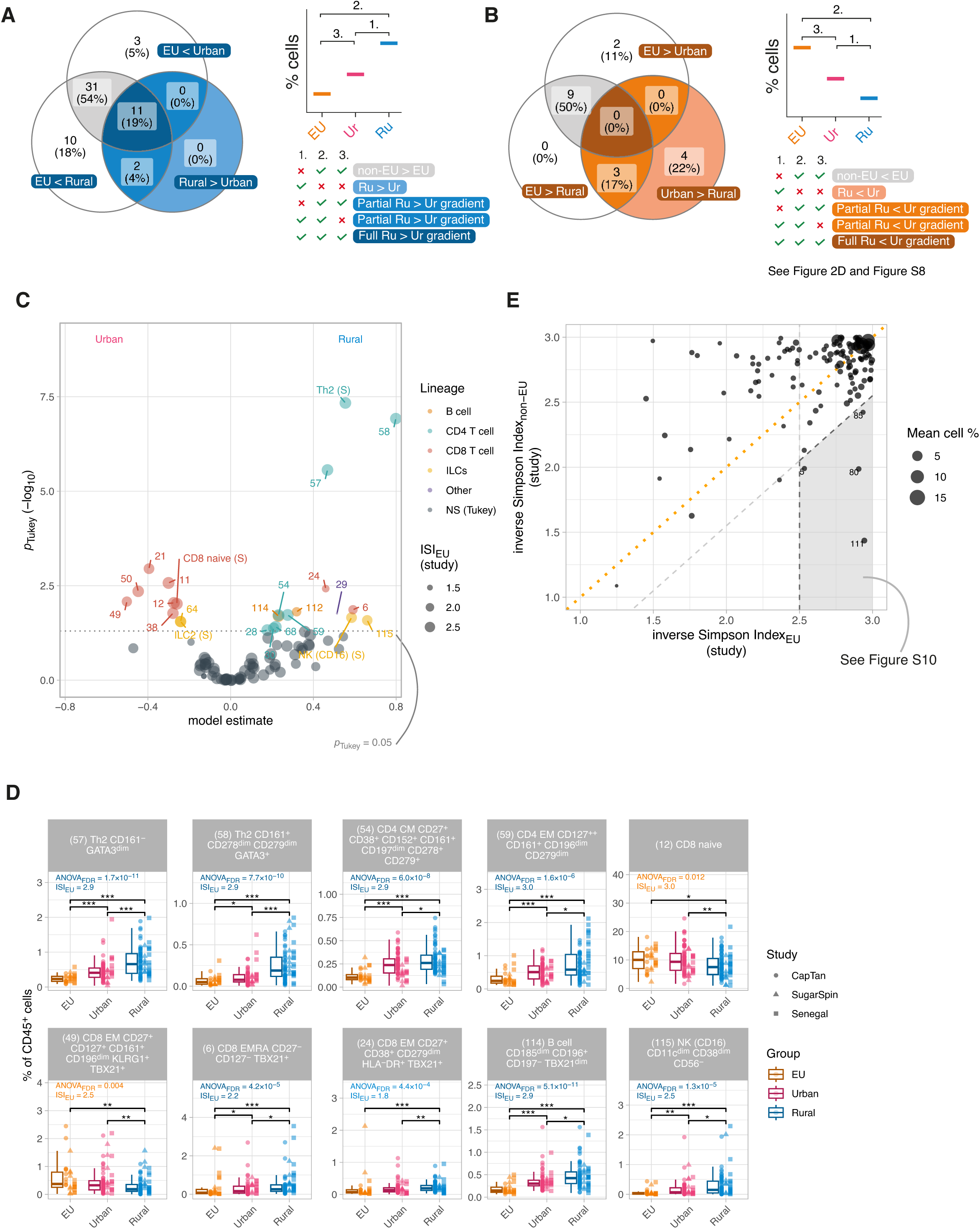
Differential cell frequencies between urban and rural-living individuals. A) and B) Venn diagrams indicating the number cell clusters that show differences in cell frequencies between 1) rural and urban-living non-Europeans, 2) rural-living individuals and Europeans and 3) urban-living non-Europeans and Europeans. Colors indicate the direction of the EU-urban-rural-gradient: A) blue, positively associated with rural-lifestyle; B) orange, negatively associated with rural-lifestyle. The strength of the EU-urban-rural-gradient was based on the number and directionality of significant pairwise comparisons (Tukey-adjusted *p*-value < 0.05; see schematic right of each Venn diagram). Significance for all comparisons was considered strongest evidence of a true rural-urban gradient (full; 1., 2. and 3.), followed by 1. and 2. or 2. and 3. (partial), and only urban-compared to rural-living individuals (Ru-Ur/limited; 1.). Significance of 2. and 3. (i.e. not 1.) was suggestive of European compared to non-European differences, rather than evidence for a rural-urban gradient and these comparisons were therefore not further considered (grey). C) Volcano plot showing differential frequency results at subset and cell cluster level. Statistical significance of pairwise comparisons was derived from a linear mixed model with cell frequency as outcome, group (EU/rural/urban) as fixed effect and sample ID and study ID as random effects. Model estimates and corresponding −log_10_-transformed Tukey-adjusted *p*-values for rural- compared to urban-living individuals were extracted using the *emmeans* R-package. Analyses were performed on *n* = 172 individuals. Each point represents a subset/cell cluster, subsets/clusters with *p_Tukey_*-values<0.05 are colored by cell lineage and labelled with subset name/cluster ID. D) Box plots showing cell frequencies for a representative selection of cell clusters (10 out of in total 20; focusing on those clusters showing a full or partial EU-rural-urban gradient; remaining clusters shown in Figure S7). Boxplots represent the 25^th^ and 75^th^ percentiles (lower and upper boundaries boxes, respectively), the median (middle horizontal line), and measurements that fall within 1.5 times the interquartile range (IQR; distance between 25^th^ and 75^th^ percentiles; whiskers). Directly right of the each boxplot, individual data points are shown. The shape and dodge of the data points corresponds with the study the samples originated from. Statistical significance of pairwise comparisons was derived from a linear mixed model with cell frequency as outcome, group (EU/rural/urban) as fixed effect and sample ID and study ID as random effects. Tukey-adjusted *p*-values for pairwise comparisons between Europeans, non-European rural- and urban-living individuals were extracted using the *emmeans* R-package. For each panel, the BH-adjusted ANOVA_FDR_-value of the differences across Europeans, non-European urban- and rural-living individuals is shown. The inverse Simpson Index in European samples (ISI_EU_) is also depicted, where higher ISI_EU_ scores indicate better integrated cell clusters. Text colour corresponds with the colors in the Venn diagrams in A). Asterisks denote statistical significance (*, *p_Tukey_* ≤ 0.05; **, *p_Tukey_* ≤ 0.01; ***, *p_Tukey_* ≤ 0.001). E) ISI_EU_ compared to ISI_non-EU_ scores. ISI_non-EU_ scores that were substantially lower (defined as >0.45 difference) compared to ISI_EU_ scores were considered possible population-specific cell clusters (only considering well integrated clusters with ISI_EU_ >2.5).

Analysis of CD4^+^ T cell clusters, showed that the T cell compartment was in a more differentiated state in rural individuals. Within this compartment, three clusters of CD161^+^ CD4 T cells (cluster 54, 59 and 68) were strongly associated with a rural lifestyle, showing a full rural > urban gradient (Figure 3D and Figure S7). Phenotypically, these clusters were all positive for CD161 and exhibited a strongly exhausted profile, expressing the immune checkpoint receptors CD279 (PD-1) and CD152 (CTLA-4). However, these clusters varied in expression of CD27, CD127, and CD196 (CCR6), possibly indicating a different degree of antigen experience and their tissue-specific homing capacity.

The most defining difference between urban- and rural-living individuals was found within the T helper (Th) 2 cell compartment. At the subset level, we found that Th2 cells, characterized by CD294 (CRTH2) expression, had higher frequencies in rural individuals compared to urban-living and European individuals (median 0.9% vs. 0.5% and 0.3% of total CD45^+^ cells, respectively; Figure S8). At the cluster level, two Th2 subpopulations were identified: GATA3^+^CD161^+^ (cluster 58) and a GATA3^dim^CD161^−^ population (cluster 57; Figure 3D). Cluster 58, which also showed increased expression of activation markers such as CD278 (ICOS), CD279 (PD-1), and KLRG1, displayed the strongest differences between rural- and urban-living individuals (Figure 3D), with smaller differences between urban-living individuals and Europeans. However, cluster 57, the less differentiated cluster of Th2, showed a more linear relationship across the full urban to rural gradient, with larger effect sizes between groups. This linear trend may be interpreted as lifetime exposure to a type 2 immunity-inducing environment.

### Urban-living individuals show higher levels of naive and CD161^+^ CD8 T cells

Within the CD8^+^ T cell compartment, we identified eight clusters that differed significantly between urban- and rural-living individuals. Two clusters were associated with rural and six were associated with urban lifestyle. The differences between rural- and urban lifestyle-associated CD8^+^ T cell clusters were best explained by their differentiation state: CD8^+^ T EM and EMRA subsets were enriched in rural, whereas naïve CD8^+^ T cells were enriched in urban-living individuals (Figure 3A-C). The CD8^+^ T cell cluster strongest associated with rural lifestyle and exhibiting a full rural > urban gradient, comprised EMRA CD8+ T cells that expressed TBX21 (Tbet), but lacked CD27 and CD127 (cluster 6; Figure 3C-D). Similarly, we identified an EM CD8 ^+^ T cell cluster (cluster 24) that expressed T-bet and CD38, together with increased expression of CD279 (PD-1), resembling the rural lifestyle–associated CD4+ T cell cluster with high PD-1 and CTLA-4 expression.

Naive CD8^+^ T cell clusters showed a different pattern, for instance a cluster of naïve CD8^+^ T cells (cluster 12) was clearly lower in rural-living individuals, but had similar cell frequencies in Europeans and urban living individuals (median 10.0% compared to 9.4% of total CD45^+^ cells, respectively; Figure 3D). In addition, we identified a cluster of CD161^bright^ CD8^+^ T cells, likely representing mucosal-associated invariant T (MAIT) cells (cluster 49). Together with cluster 50, these cells form a distinct subpopulation of CD8^+^ T cells that is largely depleted in rural-living individuals (Figure S9).

### Mature NK and B cells are enriched in rural-living individuals

In the innate part of the immune system differences between urban and rural are mostly characterized by differences between NK cells. The NK (CD16) subset, compromising of two CD16^+^ and CD56^dim/−^clusters, are highest in rural living individuals and are significant on the full rural urban scale (Figure 3D and Figure S7). More specifically, we identified a cluster (cluster 115) of CD56^−^ CD16^+^ CD11c^dim^ CD38^dim^ (IL2RB) NK cells, which were associated with rural lifestyle. In the Europeans, these mature NK cells were only found in a small abundance (median 0.02% of total CD45^+^). Within the non-European population the median expression was respectively 0.06% and 0.17% for urban and rural living individuals. In some of the rural living individuals the percentage of cluster 115 even goes above 1% of total CD45^+^ (Figure 3D). In other innate lymphoid cells, a lower frequency of ILC2s could be observed in rural living individuals, characterized by their positive expression for CD25, CD161 and CD294 (CRTH2). These cells were significantly different to the urban population, but not the European individuals (Figure S6).

Other cell subsets and clusters within the innate compartment, such as monocytes and DCs, do not seem to differ between urban and rural living individuals. However, the CyTOF panels used are limited in defining more granular innate subsets within these immune cell populations. When comparing these individuals to the Europeans, we do see clear differences. For example, the classical monocytes compartment is almost twice as large in frequency than the non-European individuals. For Europeans these cells comprise around 14.5% of total immune cells, whereas for non-Europeans this was 8.6 %. Within the B-cell compartment, we identified a cluster of CCR6^+^ B cells associated with rural compared to urban-living (cluster 114; Figure 3D).

### Several well-integrated clusters show population specific differences

To identify population-specific differences across geographical contexts (Indonesia, Tanzania, and Senegal), we compared cluster-specific ISI scores using a complementary approach. ISI values were first evaluated within European samples only, where they reflect integration quality, as each dataset contained its own European cohort. We then compared these values to ISI scores calculated for non-European samples, for which the ISI captures both integration quality and biological variation between populations. A pronounced decrease in ISI in non-Europeans relative to Europeans therefore indicates population-specific biological differences within otherwise well-integrated clusters.We identified four cell clusters demonstrating this pattern (Figure 3E). All four clusters showed significant differences between populations, with clusters of NK (CD56) CD16^+^ CD122^+^ CD161^+^ CD335^dim^ KLRG1^+^ and CD8^+^ EMRA KLRG1^dim^ specifically enriched in Indonesians. The increased frequency of CD4 EM and naive CD127^+^ CD27^+^ cells in Tanzanian and Senegalese participants compared to their Indonesian counterparts underscores significant geographical differences in immune cell distribution (Figure S10).

### The cellular immune signature can be used to robustly predict living location

To capture immune differences beyond the clusters identified by the linear mixed models, we next applied a multivariate modeling approach to predict urban versus rural living. This strategy allowed us to assess the combined predictive value of immune cell composition and functional state. Guided by the observed differences in activation and differentiation, particularly within CD4⁺ and CD8⁺ T cell compartments, we constructed an integrated immune feature set for classification.

This feature set comprised an overall immune diversity (Shannon index), relative frequencies of 26 cellular subsets, and the 80th percentile expression of eight key activation markers calculated at the subset level, yielding 235 features per individual. The use of high-quantile marker expression captured activation states while remaining robust to outliers. All features were derived from automatically annotated cell subsets, minimizing bias from manual gating or subjective cell labeling. Using this integrated immune signature, we trained machine learning models that robustly discriminated between urban and rural living within the non-European cohort.

Across nine machine learning models, seven achieved a median training accuracy greater than 0.60, indicating moderate to good performance while maintaining generalizability (Figure 4A). The best-performing model, the Lasso and Elastic-net regularized generalized linear model (glmnet), consistently showed stable training accuracy and the highest test accuracy of 0.70. Notably, the model’s feature selection reinforced key biological insights, among the top twelve most selected features, one represented a cellular subset frequency (Th2 *frequency*). Confirming the results from the generalized linear mixed models and its ability to distinguishing between urban and rural lifestyles.

**Figure 4.**
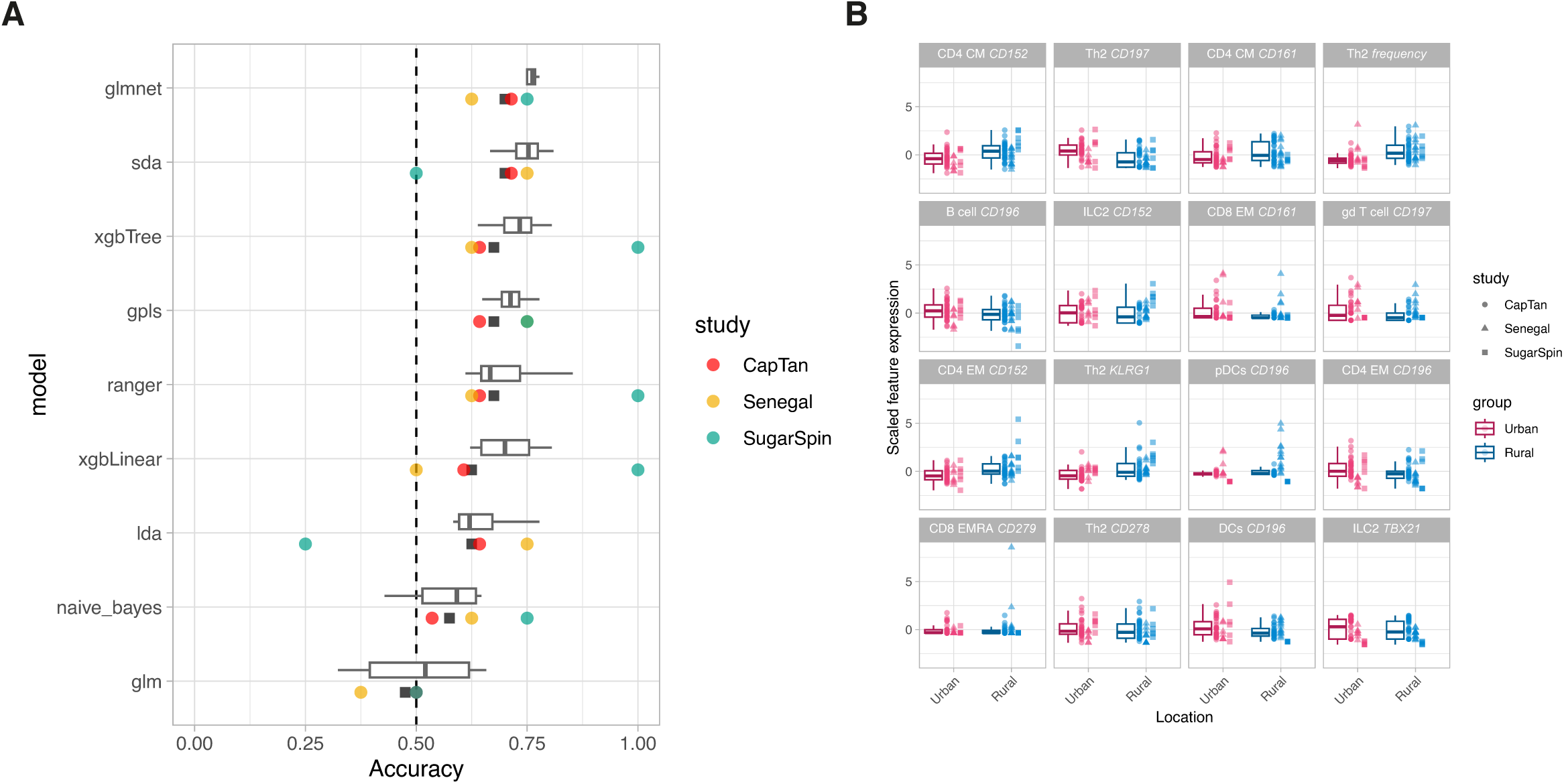
Classification models based on immune signatures are able to discriminate urban-and rural-living individuals. A) Boxplots depicting the model accuracy on the training data. All models were trained ten times with resampled train data. Dots below the respective boxplots represent the accuracy based on the test data. The black square represents the accuracy on the complete test data and colored dots indicate the study, red, yellow and green, respectively, for CapTan, Senegal and Sugarspin. Boxplots represent the 25^th^ and 75^th^ percentiles (lower and upper boundaries boxes, respectively), the median (middle horizontal line), and measurements that fall within 1.5 times the interquartile range (IQR; distance between 25^th^ and 75^th^ percentiles; whiskers). B) Boxplots of the sixteen highest contributing parameters obtained from the best performing model (glmnet).

The other eleven features were activation markers, highlighting that expression and differentiation state of immune subsets are dominant predictive factors (Figure 4B). For instance, KLRG1 expression on Th2 cells was strongly associated with rural living, whereas higher CD197 (CCR7) expression correlated with urban environments. The single most predictive feature, selected 100% of the time, was CD152 (CTLA4) expression on CD4 central memory cells, suggesting a link between rural living and sustained antigenic exposure. Additionally, CD196 (CCR6) emerged as a key marker, appearing four times among the top predictors. Elevated CCR6 expression on CD4 effector memory T cells, B cells, and dendritic cells was characteristic of an urban profile, while its expression on plasmacytoid dendritic cells was more indicative of a rural lifestyle. Together, these results suggest that differences in immune activation status and memory phenotypes, rather than cell frequencies alone, are key determinants of the immune signature associated with urban versus rural living, consistent with the patterns observed in the cluster-wise linear mixed models.

## Discussion

In this study, we used three mass cytometry datasets to study shared- and study-specific effects in immune signatures among Europeans and non-European rural- and urban-living individuals. We applied and benchmarked data-integration methods, finding strong differences in integration quality between methods. Using the top-performing integration method we were able to assess in detail cellular immune profiles associated with rural- or urban-living location shared across geography. Generally, rural-living individuals showed an immune profile that was marked by an overall highly differentiated and activated state. Using machine learning models, we were able to discriminate rural-and urban-living individuals based on these profiles with moderate to high accuracy.

In literature, several methods for batch normalization within mass cytometry datasets have been proposed, including quantile and CytoNorm normalization. CytoNorm (Van Gassen et al., 2020) performs a cluster-specific batch-adjustment for each marker by aligning identical technical replicates towards a predefined goal distribution, allowing for cell type specific marker intensity adjustments.

However, existing datasets typically lack shared technical replicates, hampering the use of this method for between-study batch-correction. Here, we showed this issue can be solved by generating an *in silico* reference sample through subsampling cells across study groups, resulting in superior integration metrics. Other methods, notably CytofIn (Lo et al., 2022), despite abrogating the need for shared technical replicates between studies, performed less well based on LISI scores. Methods applied in the single-cell sequencing field (Luecken et al., 2022), including Harmony (Korsunsky et al., 2019), also showed lower performance and typically do not scale well across 10s to 100s of millions of cells. The ability to preserve cell labels before and after integration was additionally used as a measure of data integration. Automatic cell labels were preserved at lineage level, but performance diminished at subset or cluster level, which is in line with previous work assessing cell population identification algorithms (Abdelaal et al., 2019). Nonetheless, the automatic cell labeler was a highly suitable tool for assisting in phenotyping immune cells and represented a step forward in harmonizing cell type annotations across studies. This labor-intensive work often demands profound immunological expertise and is an essential prerequisite for understanding immune cell diversity and functionality (Li et al., 2025).

Our data showed an overall relative enlargement of the memory pool within the CD4^+^ and CD8^+^ T cell lineages in non-European rural- compared to urban-living non-Europeans and Europeans, which is in line with previous findings. Among others, previous work in Senegalese urban- and rural-living participants showed higher frequencies of memory CD4^+^ T and B cells (Manurung et al., 2025; Mbow et al., 2014). We also confirm a strong Th2 polarization in rural-living individuals (de Ruiter et al., 2020) and highlight that the largest difference to urban living individuals is within the Th2 cell population that show a more differentiated phenotype, characterized by CD161 expression. These CD161^+^ GATA3^+^ Th2 cells, previously recognized as pathogenic effect Th2 (peTh2) cells, are thought to be driven by chronic antigen exposure and have an enhanced effector function (Gazzinelli-Guimaraes et al., 2024; Mitson-Salazar et al., 2016). Whereas the difference in these peTh2 cell frequencies between urban and rural-living individuals is striking, the difference between non-European and European urban-living individuals is very subtle, suggesting that indeed adopting an urban lifestyle is linked to a reduction in Th2 inducing antigenic exposures. Contrary to peTh2 cells, conventional Th2 cells (cTh2) show a very gradual increase of expression across the full urban to rural gradient, which might reflect more the lifetime exposure to Th2 inducing environment.

The increased expression of C-type lectin-like receptor CD161 in rural-living individuals was not limited to Th2 cells, but also emerged on other CD4^+^ T cells, both in the effector and central memory compartment. For example, one cluster of CD4^+^ CD127^++^ CD161^+^ was strongly associated with rural-living individuals and showing a clear trend across the full urban rural gradient. The bright expression of the interleukin-7 receptor (CD127) can be interpreted as another indication of continuous exposure to an environment rich in microorganisms and parasites (Mazzucchelli & Durum, 2007). Furthermore, CD161^+^ CD4^+^ T EM cells have previously been associated with naturally acquired immunity against malaria in endemic areas, and could be induced after repeated exposure to malaria (de Jong et al., 2021). In addition, tumor-infiltrating CD161^+^ EM CD4^+^ T cells are associated with improved survival in human papillomavirus (HPV)16-associated tumors and could be induced through therapeutic vaccination with HPV16 vaccine (Duurland et al., 2022), which might support a role for these cells in immunity. Similarly, higher CTLA-4 (CD152) was more strongly expressed in both central and effector memory CD4+ T cells in rural-living individuals, which was most apparent in our prediction model results. CTLA-4 is constitutively expressed in regulatory T cells, but also upregulated in conventional T cells after activation. This could result from repeated antigenic stimulation, which is likely to lead to immune exhaustion and, in turn, increased expression of inhibitory receptors such as CTLA-4 and PD-1 (van Dorst et al., 2024). The programmed cell death protein (PD-1) was indeed one of the other exhaustion markers that showed increased expression across the different immune lineages captured by several clusters that were increased in rural living individuals, including CD8 EM, CD4 EM, Th2 cells and Tregs.

Whereas CD161 expression on CD4^+^ T-cells appears to be a hallmark of individuals living in rural settings, the opposite is true for CD8^+^ T cells. Both from the univariate analysis and prediction models, CD161 expression on CD8 T cells is strongly associated with urban-living individuals. We identified two clusters of CD161^bright^ CD8^+^ T cells, which could be considered as mucosal-associated invariant T (MAIT) cells, but should be validated by identifying the presence of an invariant T cell receptor (Vα7.2). Both these clusters were lower in rural- compared to urban-living individuals and highest in European individuals. Abundance of MAIT cells varies widely among people and is particularly low in people at high risk of disease, including the very young, the elderly and the immunocompromised (Godfrey et al., 2019).

Within the innate immune system, we identified a cluster of CD56^−^ CD16^+^ CD11c^dim^ NK cells being elevated, with the second highest effect size, in rural living individuals. It is suggested that these cells are induced by chronic disease, and that individuals with these NK cells could respond differently to NK-mediated immunotherapies, infections or vaccines (Forconi et al., 2020). From the machine learning models we found that the expression of CD196 (CCR6) on plasmacytoid dendritic cells (pDCs) was lower in urban-living individuals, while it was higher in other dendritic cell clusters (DCs). CCR6 (CD196) and its ligand CCL20 can be related to several lung and gut disorders and play an important role in mucosal immunity (Ito et al., 2011). The observed variation in CCR6 expression across DC subsets could influence the behavior and function of these cells and may reveal mechanisms by which vaccines elicit different immune responses across populations.

This is the first study aimed at integrating mass cytometry data across healthy individuals living across the European-non-European urban-rural gradient. However, there are several limitations in our study. First, data integration will always be a balancing act between removing study-specific effects versus preserving biological signals. Therefore, even after integration, we retrieve study-specific cell clusters, which could signal either suboptimal integration or true population-specific effects. Leveraging the availability of samples from European individuals in each study, we tried to discriminate between these scenarios. Second, the addition of new data would require rerunning the normalization and clustering algorithms. However, through relatively simple adjustments, for example by defining the goal distribution as the reference of a given study or current set of studies, we could make this an expandable dataset.

Factors underlying immune variation between urban and rural lifestyle include diet (Temba et al., 2021), CMV and parasitic infections (de Ruiter et al., 2020), (early life) microbial exposures and resulting epigenetic changes, and differences in microbiota composition (Pyuza et al., 2025; Strazar et al., 2021)- and microbiome-derived metabolites (Temba et al., 2021). Importantly, these factors may vary between rural populations, although the downstream immunological effects predisposing to altered vaccine responsiveness may in fact be the same. This work underlines this idea, mainly identifying shared immunological hallmarks between genetically distinct rural populations. As such, these hallmarks may serve as targets for further prevention or intervention studies.

## Methods

### Study design

Healthy adults living in rural and urban areas (Senegal, Tanzania and Indonesia) and in urban areas in the Netherlands were included in this study.

Details on study design and in-/exclusion criteria on the study conducted in Indonesia (‘SugarSpin’) were published previously (de Ruiter et al., 2020; Tahapary et al., 2015). In short, the Indonesian samples were part of the SUGARSPIN trial, a household-based cluster-randomized double-blind trial that was conducted in three rural villages in Nangapanda, Flores Island, Indonesia. The trial was approved by the ethics committee of the Faculty of Medicine, Universitas Indonesia (FKUI) (reference no. 549/H2·F1/ETIK/2013) and filed by the ethics committee of Leiden University Medical Center (LUMC). The trial is registered as a clinical trial (reference no. ISRCTN75636394). Written informed consent was obtained from participants before the study. Age- and sex-matched samples of healthy volunteers from The Netherlands or from urban centers in Indonesia, including central Jakarta, without helminth infections were additionally included.

For the ‘Senegal’ study, details on the study were previously published (Manurung et al., 2025). In short, samples were collected from healthy Senegalese adults living in rural (Pakh and Richard-Toll) and urban (Dakar) areas and in urban areas in The Netherlands. Participants were included in the trial if they had been living in their respective residence areas for ≥10 years, preferably lifelong, were 18-40 years of age, did not have chronic diseases, and tested negative for schistosomiasis and malaria.

Samples from healthy Tanzanian adults were collected in the context of the ‘CapTan’ trial, aimed at assessing differences in immunogenicity of the yellow fever vaccine between urban and rural-living individuals. Details on the study conducted were previously published (Pyuza et al., 2024). To summarize, the study was approved by the Ethical Board of the Kilimanjaro Christian Medical University College (No. 2588) and by the Tanzania National Ethical Committee Board (NIMR/HQ/R.8a/Vol.IX/4089). Samples were collected from two rural areas (Upper Moshi and Mwanga) and two urban areas (Lower Moshi and Arusha). In addition, healthy adults living in an urban area in The Netherlands were included, which were part of the ‘TINO’- study (ClinicalTrials.gov, reference no. NCT06039527). The study was approved by the Ethics Committee of Leiden University Medical Center (NL77841.058.21).

### Parasitology

Fresh stool samples of rural Indonesian, Tanzanian and Senegalese rural-/urban-living adults were microscopically screened for soil-transmitted helminth infections (hookworms, *Ascaris lumbricoides*, *Trichuris trichiura* by Kato-Katz. Tanzanian and Senegalese adults were additionally tested for and *Schistosomiasis mansoni* (Kato-Katz). Given the low prevalence of parasitic infections in The Netherlands, Europeans were assumed negative for parasitic infections.

### Sample collection and PBMC isolation

From the participants selected for the immunology study, venous blood was collected in heparin, EDTA and dry tubes for cell isolation, full blood count (SugarSpin and Senegal only), and serology tests. From each site, peripheral blood mononuclear cells (PBMCs) were isolated from heparinised blood within four hours following blood collection, using standard Ficoll density gradient. The cells were frozen in 20%FCS/10%DMSO/RPMI freezing medium and stored in liquid nitrogen. Freezing medium was supplemented with 100 U/ml penicillin (Gibco), 100 U/ml streptomycin (Sigma-Aldrich), 1 mM pyruvate (Sigma-Aldrich), 2 mM glutamate (Sigma-Aldrich).

### Mass cytometry

#### Mass cytometry antibody staining

Antibody panels were designed to phenotype immune cells *ex vivo*. Details on antibodies used are listed in Table S3. Antibody-metal conjugates were either purchased or conjugated using a total of 100µg of purified antibody combined with the Maxpar X8 Antibody Labelling Kit (Fluidigm, San Francisco, CA, USA) according to the manufacturer’s protocol V7. The conjugated antibody was stored in 200µl of Antibody Stabilizer PBS (Candor Bioscience GmbH, Wangen, Germany) at 4°C. All antibodies were titrated on PBMC samples.

On the day of the staining, cryopreserved PBMCs were thawed with 20% FCS/2mM Mg^2+^/1:10.000 benzonase/RPMI medium (SugarSpin and Senegal: 50% FCS/RPMI medium) at 37°C and washed twice with 10% FCS/RPMI medium. For phenotyping, 3 × 10^6^ cells per sample were stained based on the Maxpar Nuclear Antigen Staining Protocol V2 (Fluidigm). First, cells were washed with Maxpar staining buffer and centrifuged for 5 min at 400g (SugarSpin and Senegal: 300g) in 5ml Eppendorf tubes.

For the Senegal and CapTan study, samples were barcoded and measured in two and seven batches, respectively. In every batch, one aliquot of PBMCs from a reference sample was included to normalize staining between batches, resulting in CN_within_-normalised data. To barcode the samples, PBMCs were resuspended in 50μl of Maxpar staining buffer, and 50μl of barcode mix targeting β2-microglobulin (B2M) was added to each individual sample in a 6-choose-3 scheme using cadmium (Cd)_106_, Cd_110_, Cd_111_, Cd_112_, Cd_114_, and Cd_116_. Samples were incubated for 30 min at room temperature and then washed with 4ml of Maxpar Staining Buffer. Cells were centrifuged for 5 min at 400g, and the supernatant was removed, resuspended and combined into 1 tube in Maxpar staining buffer.

Then, the cells were incubated with 5ml (SugarSpin and Senegal: 1ml) of 500× diluted 500µM Cell-ID Intercalator-103Rh (Fluidigm) in staining buffer at room temperature for 15 min to identify dead cells. After washing with staining buffer, cells were incubated with 20µl (SugarSpin and Senegal: 5µl) of Human TruStain FcX Fc receptor blocking solution (BioLegend) and 130µl (SugarSpin and Senegal: 45µl) of staining buffer at room temperature for 5 min (SugarSpin and Senegal: 10 min).

Then, 150µl (SugarSpin and Senegal: 50µl) of freshly prepared surface antibody cocktail was added and incubated at room temperature for another 30 min (SugarSpin and Senegal: 45 min). Subsequently, cells were washed twice with staining buffer and fixed with 1.6% PFA in 5ml PBS and incubated for 10 min (CapTan only). Cells were spun 800g for 7 min, supernatant was removed after which cells were fixed and permeabilized using the eBioscience Foxp3/Transcription Factor Staining Buffer Set (eBioscience, cat. 00-5523-00). After cells were incubated with 5ml (SugarSpin and Senegal: 1ml) of the freshly prepared Fix/Perm working solution (prepared according to the manufacturer’s instructions) for 45 min, cells were spun and washed twice with 1× permeabilization buffer at 800g for 7 min (SugarSpin and Senegal: 5 min). Next, cells were resuspended in 130µl of 1× permeabilization buffer and incubated for 5 min with Human TruStain FcX Fc receptor blocking solution (CapTan only). Then, 150µl (SugarSpin: 50µl) of the intranuclear antibody cocktail was added and incubated for 30 min at room temperature. Following incubation, cells were washed once with 1× permeabilization buffer and once (SugarSpin and Senegal: twice) with staining buffer before fixation with 1.6% PFA in 5mL PBS and incubated for 10 min (CapTan only). Cells were spun and stained with 5ml (SugarSpin and Senegal: 1ml) of 1000× diluted 125 µM Cell-ID Intercalator-Ir (Fluidigm) in Maxpar Fix and Perm Buffer (Fluidigm) at room temperature for 1h (SugarSpin and Senegal: at 4°C overnight) to stain all cells. After two (SugarSpin and Senegal: three) washes with staining buffer and centrifugation at 800g, cells were resuspended in RPMI 20% FCS 10% DMSO and stored at −80°C until acquisition. For the SugarSpin and Senegal study, cells were stored as a pellet at 4°C and measured within 2 days.

#### Mass cytometry antibody acquisition

For the Senegal and CapTan study, cells were thawed on the day of acquisition in RPMI 50% FCS, spun and washed twice with Maxpar staining buffer. Samples were measured with a Helios mass cytometer (Fluidigm), which was automatically tuned according to Fluidigm’s recommendations. Before measuring, cells were counted, washed with Milli-Q water, passed over a cell strainer, and brought to a concentration of 1.0 × 10^6^ cells/ml with 10% EQ Four Element Calibration Beads (Fluidigm) in Milli-Q water. Mass cytometry data were acquired and analyzed on-the-fly using dual-count mode and noise-reduction on. Next to channels to detect antibodies, channels for intercalators (_103_Rh, _191_Ir, and _193_Ir), calibration beads (_140_Ce, _151_Eu, _153_Eu, _165_Ho, and _175_Lu), and background/contamination (_133_Cs, _138_Ba, and _206_Pb) were acquired. After data acquisition, the mass bead signal was used to normalize the short-term signal fluctuations with the reference EQ passport P13H2302 during the course of each experiment. When applicable, normalized FCS files were concatenated using Helios software without removing the beads.

### Data analysis

All data preprocessing and statistics were performed in R v4.2.2 and RStudio Server v2022.03.999. All *p*-values were corrected for multiple testing using to the Benjamini-Hochberg procedure (and referred to as *q*-values). *q*-values < 0.05 were considered statistically significant.

#### Data preprocessing

First, raw (bead-normalized) data were preprocessed through a unified pipeline. Briefly, the pipeline included automated gating using Gaussian parameters (*CyTOFClean* R-package; v1.03beta; https://github.com/JimboMahoney/cytofclean) and automated quality control for signal stability over acquisition time (*PeacoQC* v1.9.3 R-package (Emmaneel et al., 2022); parameters recommended for mass cytometry data). Next, automatic gating was applied to select for intact/DNA^+^-(_191_Ir and _193_Ir channels), CD45^+^- (_89_Y), live cells (live/dead staining) (*openCyto* v2.10.1 R-package). All automatically set gates were manually inspected. Samples were compensated (only CapTan study) and debarcoded (CapTan and Senegal studies; *CATALYST* v1.22.0 R-package). Resulting data were converted from .FCS-files into a SingleCellExperiment-object (‘raw’). Data were transformed using a hyperbolic arcsinh-transformation with a cofactor of 5 for downstream processing, unless otherwise specified.

Next, reference samples collected from healthy European adults included in each individual batch (CapTan and Senegal studies) were used to train a CytoNorm-model (*CytoNorm* v0.0.17 R-package; CytoNorm.train-function; nQ = 101; goal = ‘mean’; *k* = 5). The trained model was applied to all samples, adjusting for batch effects within studies (CN_within_; CytoNorm.normalize-function). The SugarSpin-study was not acquired in batches and therefore was added to the CN_within_-dataset without further processing of the ‘raw’ data.

#### Data integration methods

When merging mass cytometry data, appropriately adjusting for batch effects, within and between studies, is a major challenge. Batch effects stem from differences in sensitivity across cytometers, metal sensitivity and oxidation, variability in antibodies and channel spillover (Rybakowska et al., 2021). These effects can be minimized by using standardized experimental protocols, reducing sources of technical variation. Furthermore, identical technical replicates are typically included to correct variation in marker intensities between batches within studies based on a goal distribution (Van Gassen et al., 2020). However, existing datasets generally lack identical replicates that allow adjustment of batch effects between datasets collected over time.

We here assessed four methods to adjust between-study batch effects, including CytofIn (Lo et al., 2022), Harmony (Korsunsky et al., 2019), quantile normalization and CytoNorm (Van Gassen et al., 2020). Two-step normalization approaches were not considered. For CytofIn (Lo et al., 2022) (mode = ‘meanshift’; default), the same reference samples as used to generate the CN_within_-normalized data were used. For the SugarSpin study, one randomly chosen European sample was considered the reference for this study. For quantile and CytoNorm (between studies; CN_between_), *in silico* reference samples were generated by subsampling 1M cells per study, across study groups. Reference samples were inspected using t-SNE visualizations. Next, quantile and CytoNorm were run with similar settings to the CN_within_-normalization (nQ = 101; goal = ‘mean’; *k* = 5), except that metaclustering performed using the metaClustering_hclust-function. All cells with arcsinh(5)-transformed expression values below zero or above eight were removed.

#### Cell clustering

Study-specific CN_within_-normalized and integrated CN_between_-normalised datasets were subjected to flowSOM-clustering (15 × 15 hexagonal grid; rlen=100; *kohonen* v3.0.11 R-package), followed by metaclustering at various values of *k* (total number of clusters), using the *ConsensusClusterPlus* v1.62 R-package (reps = 100, distance = ‘euclidean’). The clustering map was trained on 100k cells per sample, the remaining cells were mapped to the trained map (*predict.kohonen*-function). For choice of the optimal value for *k,* see below.

#### Diversity measures

Immune profile diversity measures (richness, Shannon and Simpson indices) were calculated using the *vegan* R-package v2.6-4. Richness was defined as the number of cluster present at >0.1% of CD45^+^-cells. Statistical significance was assessed using a linear model with group as predictor, while adjusting for study. Estimates and *p*-values for each pairwise comparison between groups (e.g. urban vs rural) were extracted using the *emmeans* v1.8.5 R-package.

#### Automated cell cluster annotations

Cell clusters before (CN_within_-normalized data) and after (CN_between_-normalised data) were subsequently subjected to an automated cell labeling algorithm. Briefly, for each given marker, the algorithm assesses the distribution of that marker across all cell clusters. Marker distributions are then clustered using hierarchical clustering to define two groups: a ‘positive’ and a ‘negative’ group of marker distributions. Subsequently, for each distribution we check how the median expression relates to both the positive and negative groups of distributions. If the median is higher than the 5^th^ percentile of the positive group, the cluster is deemed positive for that marker. If the median is lower than the 95^th^ percentile of the negative group, the cluster is deemed negative for that marker. If neither of these conditions applies, then the marker is considered neutral for that cluster. In this manner we define for each cluster whether a marker is positive, negative or neutral (i.e. not clearly positive or negative). To evaluate the performance of the automated cell labeler, post-integration cell labels were compared to pre-integration labels (considered ‘ground truth’). To measure the accuracy of the labelling, the F1 score (harmonic mean of the precision and recall) was computed, for every cell population (*yardstick*

R-package; f_meas-function; average = ‘binary’). The overall F1 score per dataset were computed as the median of F1 scores across cell populations (excluding populations labeled as ‘missing’ or ‘other’).

#### Dataset-level quantification of data integration

To assess integration, local inverse Simpson index (LISI)-scores were calculated based on 50k cells per study (150k cells total) using default parameters (perplexity = 30; neighbourhood size = 90 cells) (Korsunsky et al., 2019). LISI-scores were calculated to assess whether clusters of cells are well-mixed across studies. If cells are well-mixed, LISI-scores are near the total number of studies in our analysis (i.e. 3). Second, we used *k*-nearest-neighbor batch-effect test (kBET (Buttner et al., 2019)) to quantify batch effects. Briefly, kBET evaluates the accordance a the local fraction of cells in a given batch compared to the global fraction of cells in that batch. Cells are considered well matched if the batch label distribution of local fractions are similar to the global fraction. To estimate label distributions in local fractions, the algorithm creates *k*-nearest neighbour matrix (sample size; 900 cells) and choses 10% of the cells to check the batch label distribution in its neighbourhood (with neighborhoodsize *k0*, ranging between 1 to sample size). If the (observed) local batch label distribution in this subset of cells is sufficiently similar to the global batch label distribution, the χ^2^-test does not reject the null hypothesis (so all batches are considered well-mixed). The subsampling is performed over 100 repeats for each neighborhood size.

#### Subset-level quantification of data integration

To quantify data integration across larger subsets of cells (i.e. cluster, subset or lineage), we calculated a (global) inverse Simpson index (ISI)-score across all cells belonging to a given subset. Here, ISI-scores differ from LISI-scores, as the latter are calculated across local neighborhood of a set of cells. Despite that, ISI-scores significantly correlated with mean LISI scores for each cluster (Spearman’s rank correlation 0.43; *p-*value = 2.8 × 10^−11^), indicating that the global ISI-scores adequately captured local mixing of studies. ISI-scores were calculated across European samples, as these are expected to show no or very limited variation between studies. Therefore, any variation quantified using (L)ISI-scores is likely the result of batch-effect and not biological variation. These scores were compared to ISI-scores based on non-European samples; relatively low scores (arbitrary minimal difference >0.45) in non-Europeans compared to Europeans could indicate population-specific effects in well-integrated cell clusters (ISI_EU_ > 2.5).

#### Statistics

PERMANOVA-tests were used to assess the overall effect of study/group on the immune cell frequency data matrix (univariable tests, method = ‘euclidean’, 9,999 permutations; *vegan* R-package v2.6-4). The statistical significance of the effect of group was assessed with permutations restricted within study strata where appropriate.

Principal component analysis (PCA) was performed on cell cluster frequencies of all individuals (*prcomp* R-function). Cell frequencies were centered and scaled to have unit variance.

For visualization, cells were embedded in a two dimensional t-distributed Stochastic Neighbor Embedding (t-SNE) map using the FIt-SNE algorithm v1.2.1 (Linderman et al., 2019). FIt-SNE was performed on a downsampled dataset including 50,000 cells per study (max_iter = 1,000; learning rate = *n* cells/12; perplexity = *n* cells/100).

To compare the abundance of cell clusters between urban Europeans and rural and urban living non-Europeans we employed a generalized linear mixed model (binomial = ‘family’; link = ‘logit’; *lme4* R-package v1.1-31). The frequency for each identified lineage, subset or cluster (divided by total CD45^+^ cells) was considered the dependent variable, ‘group’ was a fixed explanatory variable and ‘study ID’ and ‘sample ID’ were included as random intercepts. ‘sample ID’ was included as random effect to deal with any under- or overdispersion due to the binomial nature of the model. This ‘full’ model was compared to a ‘reduced’ model from which ‘group’ was removed to assess (the significance of) its effect (ANOVA). Resulting *p*-values were corrected for multiple testing (simultaneously accounting for lineage-, subset- and cluster-level analyses). Following, estimates and *p*-values for each pairwise comparison between groups (e.g. urban vs rural) were extracted using the *emmeans* v1.8.5 R-package. Statistical significance of these pairwise comparisons was assessed using the Tukey *post hoc* test. To optimize the visualization of differences between groups using boxplots, we performed median absolute deviation (MAD)-filtering, not showing samples with cluster/subset frequencies > 25 × MAD.

#### Rural-urban classification

As a proof-of-principle, we used a wide range of machine learning models to investigate if we could discriminate urban- from rural-living individuals based solely on cellular immunological signatures. Modeling was performed on subset-level cell frequencies and the 80^th^ percentile of marker intensity values of activation markers (defined as CD152 [CTLA-4], CD161, CD196 [CCR6], CD197 [CCR7], CD278 [ICOS], CD279 [PD1], TBX21 [Tbet] and KLRG1) for each subset, all features were scaled to unit variance. Data was split in a train (70% of the data) and test data partition (*caret* v7.0 R-package), partitioning of data was performed by splitting the studies in equal proportion to minimize bias towards a specific study. Automated model tuning was performed on training data using 5-fold cross-validation over 10 repeats (*caretEnsemble* v4.0.1 R-package). Model performance was compared between the following models: shrinkage discriminant analysis (sda), linear discriminant analysis (lda), elastic-net regularized generalized linear model (glmnet), random forest (ranger), eXtreme gradient boosting (xgbLinear, xgbTree), least squares support vector machine (lssvmRadial), Naïve Bayes (naive_bayes), generalized linear model (glm) and generalized partial least squares (gpls).

## Acknowledgements

This study is part of the EDCTP2 program supported by the European Union and by grants from the Dutch Research Organization (NWO) through the Spinoza prize awarded to Maria Yazdanbakhsh, the European Research Council (ERC) via the ERC Advanced Grant ‘REVERSE’ awarded to Maria Yazdanbakhsh (Grant No: 101055179), the LUMC Excellent Student Fellowship awarded to Marloes M.A.R. van Dorst and the LUMC Global PhD Fellowship awarded to Jeremia J. Pyuza. We would to like acknowledge all clinical and research staff at study sites in Tanzania, Senegal and Indonesia who helped to make this study possible. We would also like to acknowledge the LUMC core facility for providing mass cytometry services. Finally, we would like to thank all volunteers who participated in this study.

## Authors contributions

Conceptualization, K.A.S., W.A.A.d.S.P., M.Y.; Methodology, J.J.P., M.M.A.R.v.D., M.K., Y.C.M.K.; Data Curation, K.A.S., W.A.A.d.S.P.; Formal Analysis, K.A.S., W.A.A.d.S.P., M.M., A.M., M.C., S.P.J.; Investigation, J.J.P., M.M.A.R.v.D., D.LT., T.S., J.P., M.M.; Writing - Original Draft, K.A.S., W.A.A.d.S.P.; Review & Editing, all authors; Supervision, A.M., S.P.J., W.A.A.d.S.P., M.Y.; Funding Acquisition, M.Y. All authors read and approved the final manuscript.

## Competing interest

The authors declare that they have no competing interest.

## Data and materials availability

All data associated with this study are present and available upon request.

## Supplementary figures

**Figure S1 | Data preprocessing and integration pipeline.** Flowchart showing the data preprocessing, quality control and integration methods applied.

**Figure S2 | kBET mean rejection rates depend by neighborhood size.** The dashed vertical lines indicate the optimal neighborhood size for batch-effect detection, that is, where the rejection rate is maximal. Shaded areas represent the 95th percentile of *n* = 100 repeated kBET runs. In each run, the number of tested neighborhoods was 10% of the sample size (i.e., 90 cells).

**Figure S3 | Integration metrics to assess clustering depth.** A) Median inverse Simpson index (within European samples; study) across all cell subsets (clusters, codes, subsets or lineages). Higher values indicate higher diversity of study-labels within a given group of cells, suggesting better integration. The optimal number of clusters was *k =* 125 (first local peak in median ISI across clusters with *k* ≥100). B) Boxplots of inverse Simpson indices (within European samples; study) for cell clusters (for a selection of *k*-values), codes, subsets and lineages. Boxplots represent the 25^th^ and 75^th^ percentiles (lower and upper boundaries boxes, respectively), the median (middle horizontal line), and measurements that fall within 1.5 times the interquartile range (IQR; distance between 25^th^ and 75^th^ percentiles; whiskers). C) and D) Stacked bar plots showing inverse Simpson index categories, indicating low (≤ 2.2), moderate (2.2 – 2.5), high (2.5-2.8) and very high (>2.8) indices. Percentages were calculated by dividing the number of cell subsets (i.e. cell clusters, codes, subsets or lineages) within a given inverse Simpson index category by the total number of cell subsets (C). In D) we additionally normalized for the number of cells within each cell subset. Here, a considerable drop in the percentage of cells allocated to a cell cluster with ISI-score >2.8 can be appreciated.

**Figure S4 | Heatmap showing median marker expression for each cluster.** Clusters were based on SOM and hierarchical clustering. Each tile depicts the median expression of a given marker (rows) for a specific cluster (columns). The heatmap is stratified based on cell lineage.

**Figure S5 | Highest cell cluster diversity in rural-living individuals compared to urban-living and European individuals.** Richness (number of cell clusters present at >0.1% of CD45^+^-cells), Shannon and Simpson diversity measures by A) group and B) group, stratified by lineage (CD4^+^ or CD8^+^ T cells). Diversity measures were calculated based on rarefied data, subsampling each sample to a ‘depth’ of 70,000 cells. Boxplots represent the 25th and 75th percentiles (lower and upper boundaries of boxes, respectively), the median (middle horizontal line), and measurements that fall within 1.5 times the interquartile range (IQR; distance between 25th and 75th percentiles; whiskers). Statistical significance was assessed using linear models with group and study as fixed effects. Statistical significance of differences between each group was assessed using the *emmeans()*-function. P-values were corrected for multiple testing using the Benjamini-Hochberg method. Asterisks denote statistical significance (*, q ≤ 0.05; **, q ≤ 0.01; ***, q ≤ 0.001).

**Figure S6 | Subset frequencies between Europeans, rural- and urban-living individuals.** Box plots showing subset frequencies (all detected subsets shown). Boxplots represent the 25^th^ and 75^th^ percentiles (lower and upper boundaries boxes, respectively), the median (middle horizontal line), and measurements that fall within 1.5 times the interquartile range (IQR; distance between 25^th^ and 75^th^ percentiles; whiskers). Directly right of the each boxplot, individual data points are shown. The shape and dodge of the data points corresponds with the study the samples originated from. Statistical significance of pairwise comparisons was derived from a linear mixed model with cell frequency as outcome, group (EU/rural/urban) as fixed effect and sample ID and study ID as random effects. Tukey-adjusted *p*-values for pairwise comparisons between Europeans, non-European rural- and urban-living individuals were extracted using the *emmeans* R-package. For each panel, the BH-adjusted ANOVA_FDR_-value of the differences across Europeans, non-European urban- and rural-living individuals is shown. The inverse Simpson Index in European samples (ISI_EU_) is also depicted, where higher ISI_EU_ scores indicate better integrated subsets. Analyses were performed on *n* = 172 individuals. Asterisks denote statistical significance (*, *p_Tukey_* ≤ 0.05; **, *p_Tukey_* ≤ 0.01; ***, *p_Tukey_* ≤ 0.001).

**Figure S7 | Differential cell frequencies between rural- and urban-living individuals.** Boxplots showing cell frequencies for cell clusters showing significant differences between rural- and urban-living individuals (remaining 10/20 clusters; 10/20 depicted in Figure 3D; *p_Tukey_*-value<0.05 for at least one comparison; see legend Figure 3D).

**Figure S8 | Th2 cell marker expression.** A-C) t-distributed Stochastic Neighbor Embedding (t-SNE) visualization Th2 cells (clusters 57 and 58; *n* = 1k random cells/subject). A) Global location of CD161^+^ and GATA^+^ Th2 cells. B) t-SNE embeddings were colored according to marker expression of CD294 (CRTH2), GATA3, CD161, KLRG1 (MAFA) and CD297 (PD-1) C) cluster allocation (based on global clustering across all cells). D) Density features of Th2 cells. E) Distribution of marker expression of CD294 (CRTH2), GATA3, CD161, KLRG1 (MAFA) and CD297 (PD-1) for each Th2 cluster (57 and 58). Area under the ridges was colored according to marker expression quartile.

**Figure S9 | Lower rates of naïve CD8+ T cells in rural-living non-Europeans.** A-B and D) t-distributed Stochastic Neighbor Embedding (t-SNE) visualization CD8^+^ T cells (based on cluster annotation; *n* = 26 clusters; *n* = 1k random cells/subject). A) Global location of CD8^+^ naïve, effector memory (EM), effector memory T cells that re-express CD45RA (EMRA), CD161^+^ EM, and central memory (CM) cells. B) t-SNE colored according to cluster allocation (based on global clustering across all cells). D) t-SNE colored according to marker expression of CD45RA, CD45RO, CD197 (CCR7), CD127 (IL-7RA), CD161, KLRGI (MAFA), CD279 (PD-1), Tbet and HLA-DR. C) Density features of CD8^+^ T cells. E) Distribution of marker expression of CD279 (PD-1), KLRG1 (MAFA), Tbet, CD161 and HLA-DR for each CD8^+^ T cell cluster. Area under the ridges was colored according to marker expression quartile.

**Figure S10 | Differential cell frequencies of well-integrated, study-specific cell clusters.** Box plots showing cell cluster frequencies showing significant differences between studies, stratified between non-Europeans (non-EU) and Europeans (EU). The selection of these clusters was based on Figure 3E. Boxplots represent the 25^th^ and 75^th^ percentiles (lower and upper boundaries boxes, respectively), the median (middle horizontal line), and measurements that fall within 1.5 times the interquartile range (IQR; distance between 25^th^ and 75^th^ percentiles; whiskers). Directly right of the each boxplot, individual data points are shown. The shape and dodge of the data points corresponds with the study the samples originated from. Statistical significance of pairwise comparisons was derived from a linear mixed model with cell frequency as outcome, group (EU/rural/urban) as fixed effect and sample ID and study ID as random effects. Tukey-adjusted *p*-values for pairwise comparisons between Europeans, non-European rural- and urban-living individuals were extracted using the *emmeans* R-package. For each panel, the BH-adjusted ANOVA_FDR_-value of the differences across Europeans, non-European urban- and rural-living individuals is shown. The inverse Simpson Index in European samples (ISI_EU_) is also depicted, where higher ISI_EU_ scores indicate better integrated cell clusters. Analyses were performed on *n* = 172 individuals. Asterisks denote statistical significance (*, *p_Tukey_* ≤ 0.05; **, *p_Tukey_* ≤ 0.01; ***, *p_Tukey_* ≤ 0.001).

## Supplementary tables

**Table S1 | Dataset metadata.** Dataset metadata for CapTan, Senegal and SugarSpin studies.

**Table S2 | Baseline characteristics across study groups.** EU, European; Urban, urban-living non-Europeans; Rural, rural-living non-Europeans. n (%) (categorical variables) or median (IQR) (continuous variables) shown, ^a^ Kruskal-Wallis rank sum test; ^b^ Pearson’s Chi-squared test.

**Table S3 | Mass cytometry antibody panel.** CCR, C-C chemokine receptor. CD, cluster of differentiation. CRTH2, prostaglandin D2 receptor 2. CXCR, CXC chemokine receptor. HLA-DR, human leukocyte antigen-D-related. IL-2R, interleukin-2 receptor. IL-7Rα, interleukin-7 receptor α. KLRG1, killer cell lectin-like receptor subfamily G member 1. MAFA, mast cell function-associated antigen. PD-1, programmed cell death protein 1. TCR, T cell receptor. RORγt, related orphan receptor gamma t.

**Table S4 | ‘Rules’ automated cell labeller.** For each lineage and subset, positive (pos), negative (neg), not positive (i.e. negative/neutral; not_pos), not negative (i.e. positive/neutral; not_neg) markers are shown. For each cell cluster, markers were classified as ‘positive’, ‘negative’ or ‘neutral’ by comparing marker intensity distribution of that cluster to the marker intensity distribution in all other cell clusters.

**Table S5 | Cell annotations.** Annotation of each cluster separated on lineage, subset and cluster level. Both the annotation of the automatic cell labeller and the revised annotation is included (manual).

## Notes

### Competing Interest Statement

The authors have declared no competing interest.

### Summary of Updates

Manuscript adapted to improve readability and consistency

