## Supplementary figures and images for "Mass cytometry data integration methods reveal rural-urban gradient of immune profiles across geography"

### Supplemental Figure 1

#### Figure S1

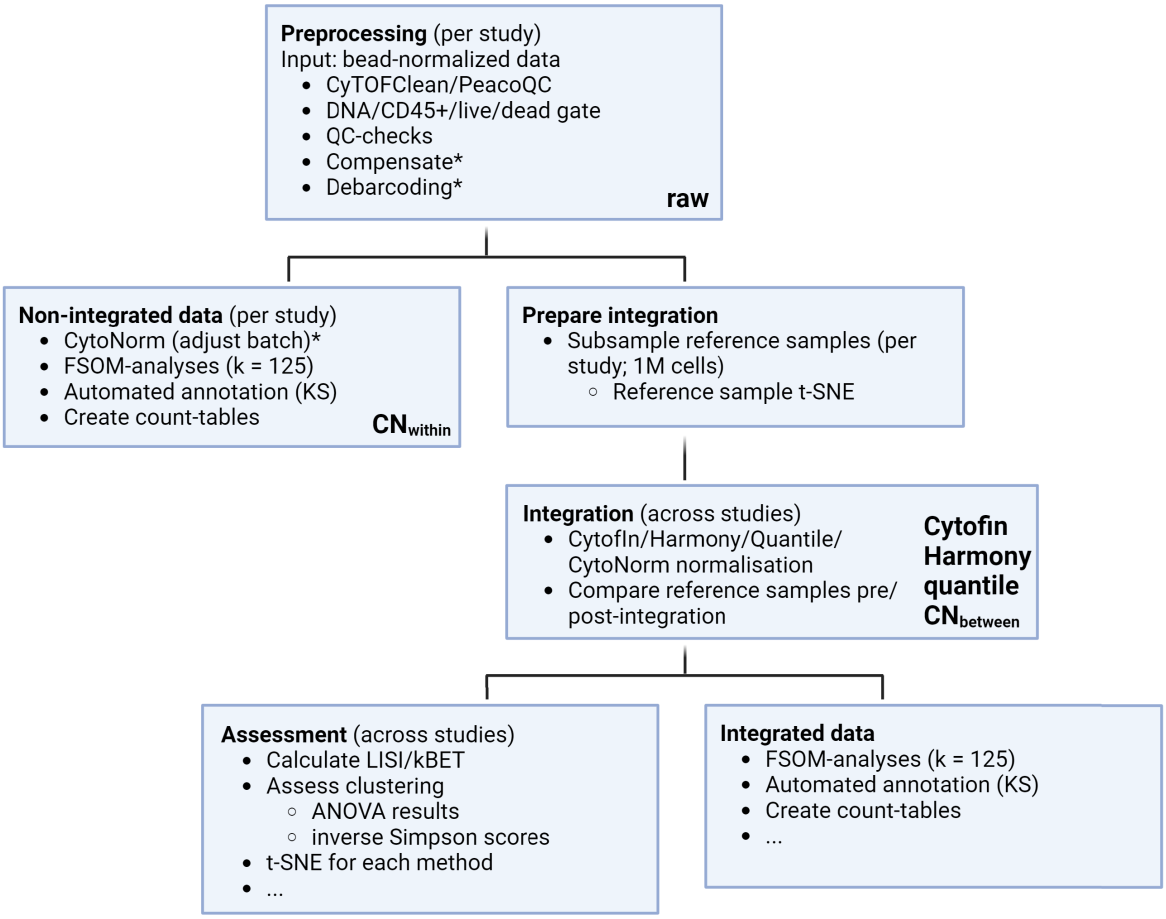


*Only for Senegal (2 batches)/CapTan (7 batches)

### Supplemental Figure 2

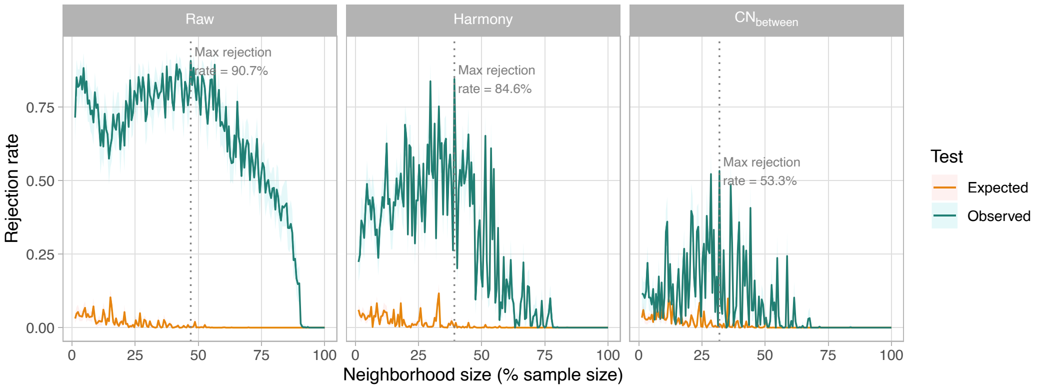

### Supplemental Figure 3

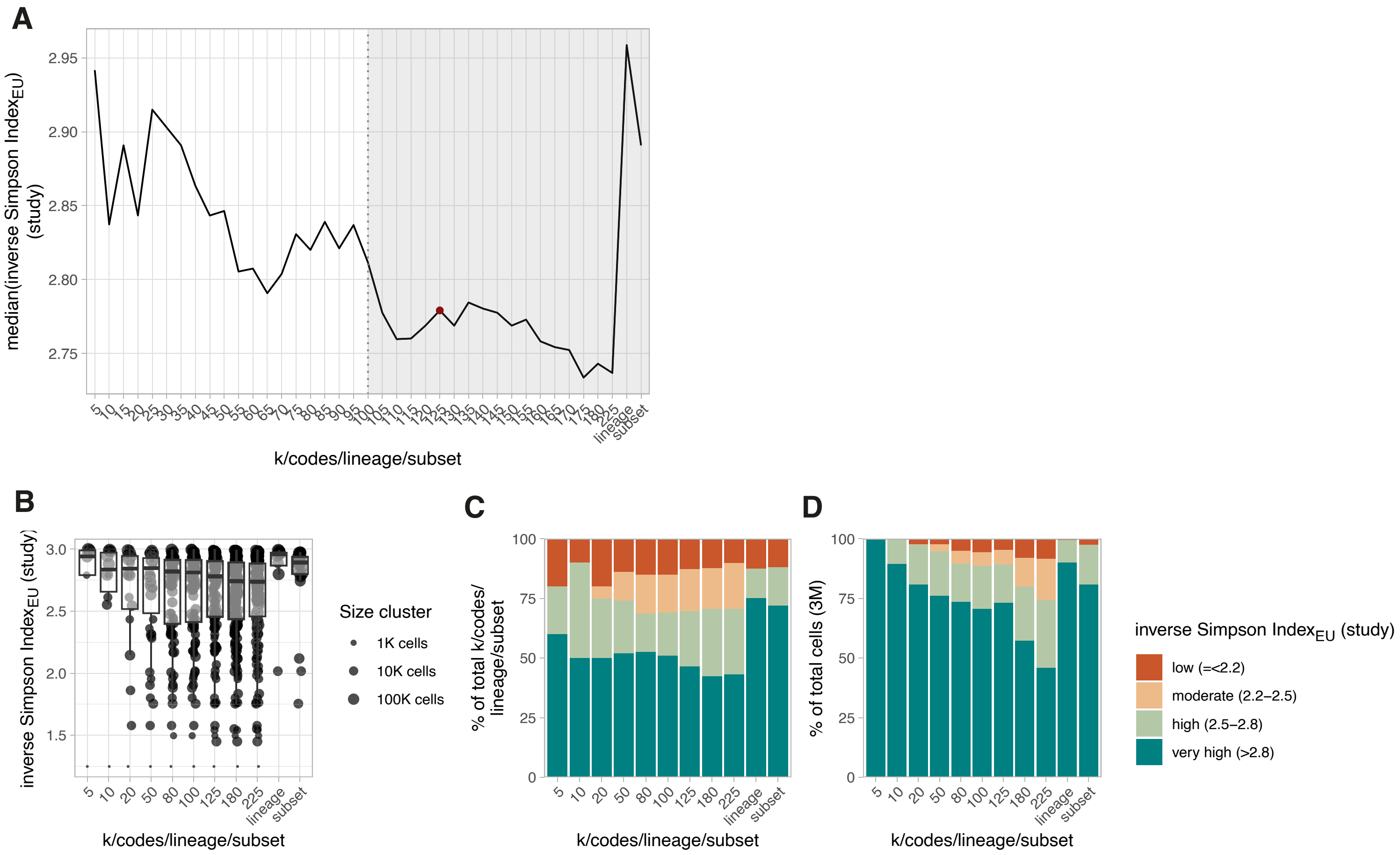

### Supplemental figure 4

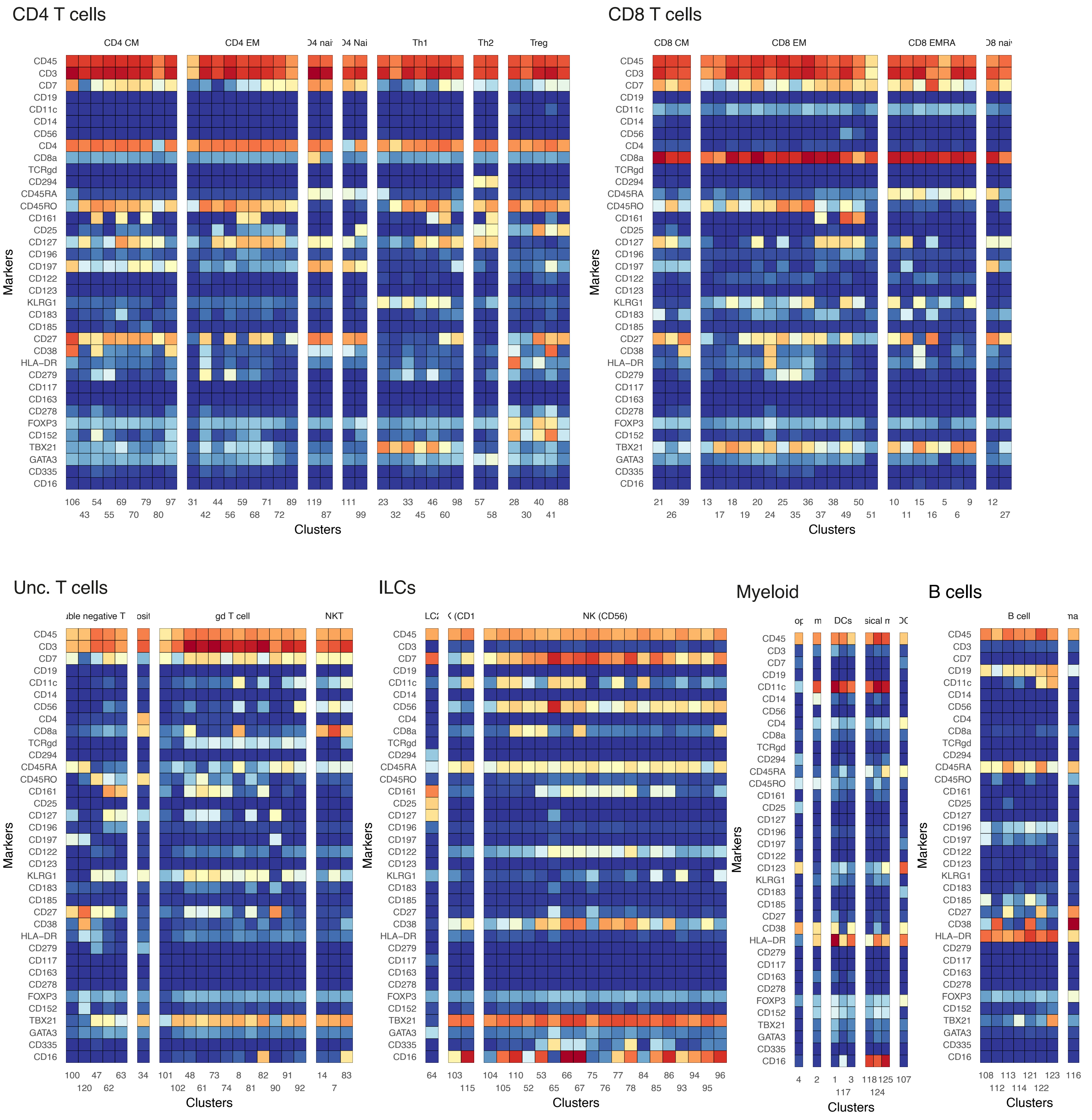

### Supplemental Figure 5

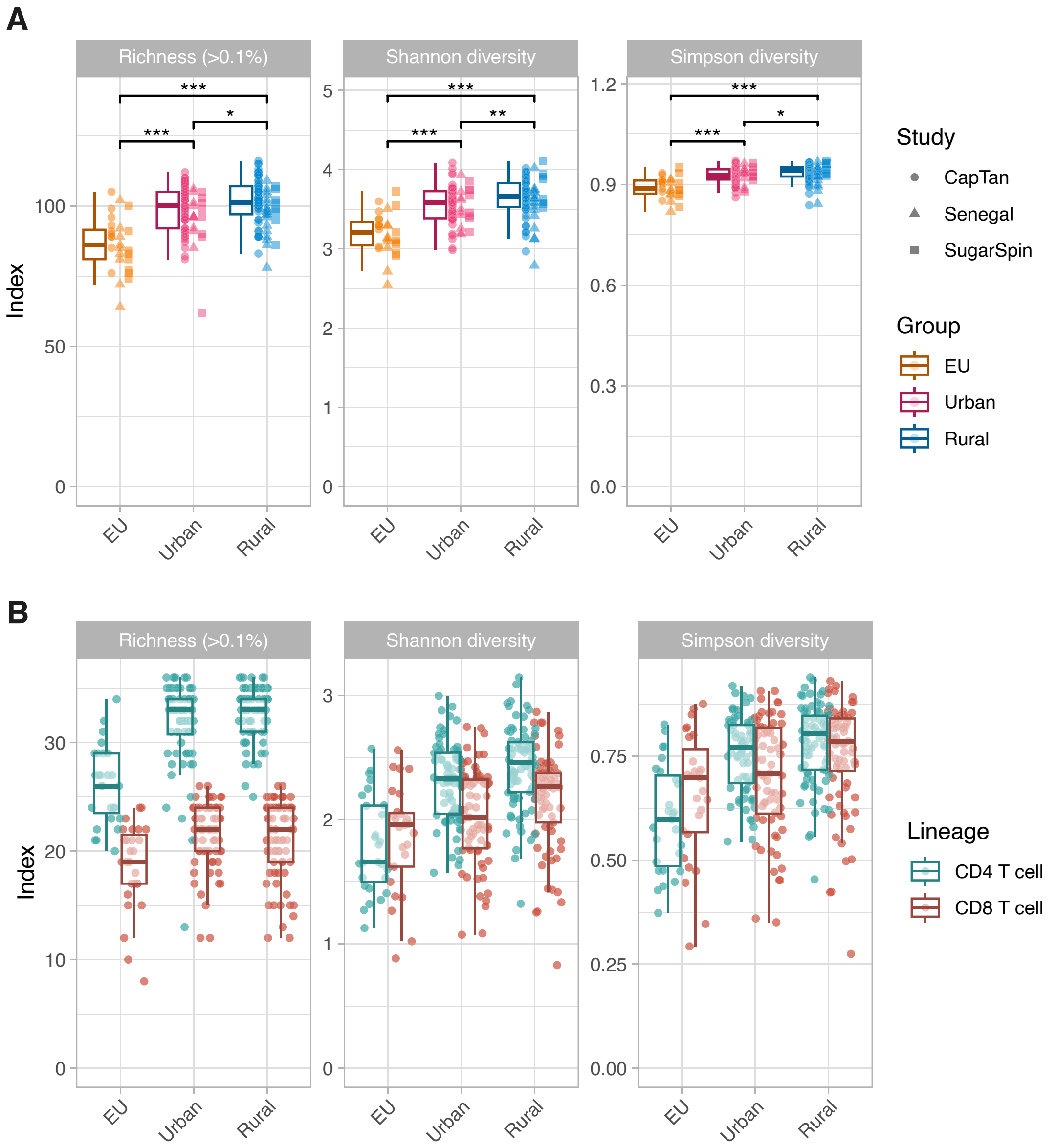

### Supplemental Figure 6

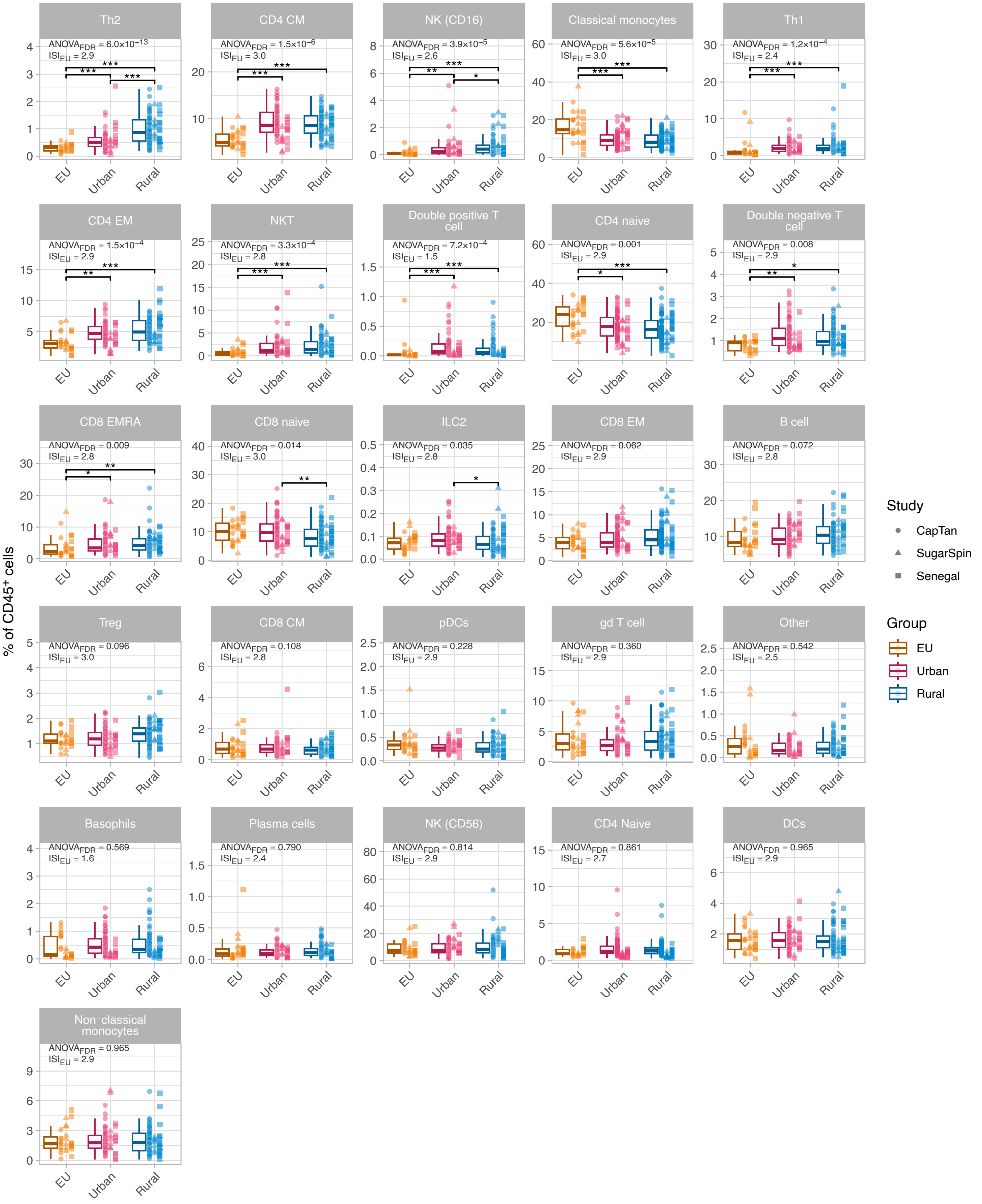

### Supplemental Figure 7

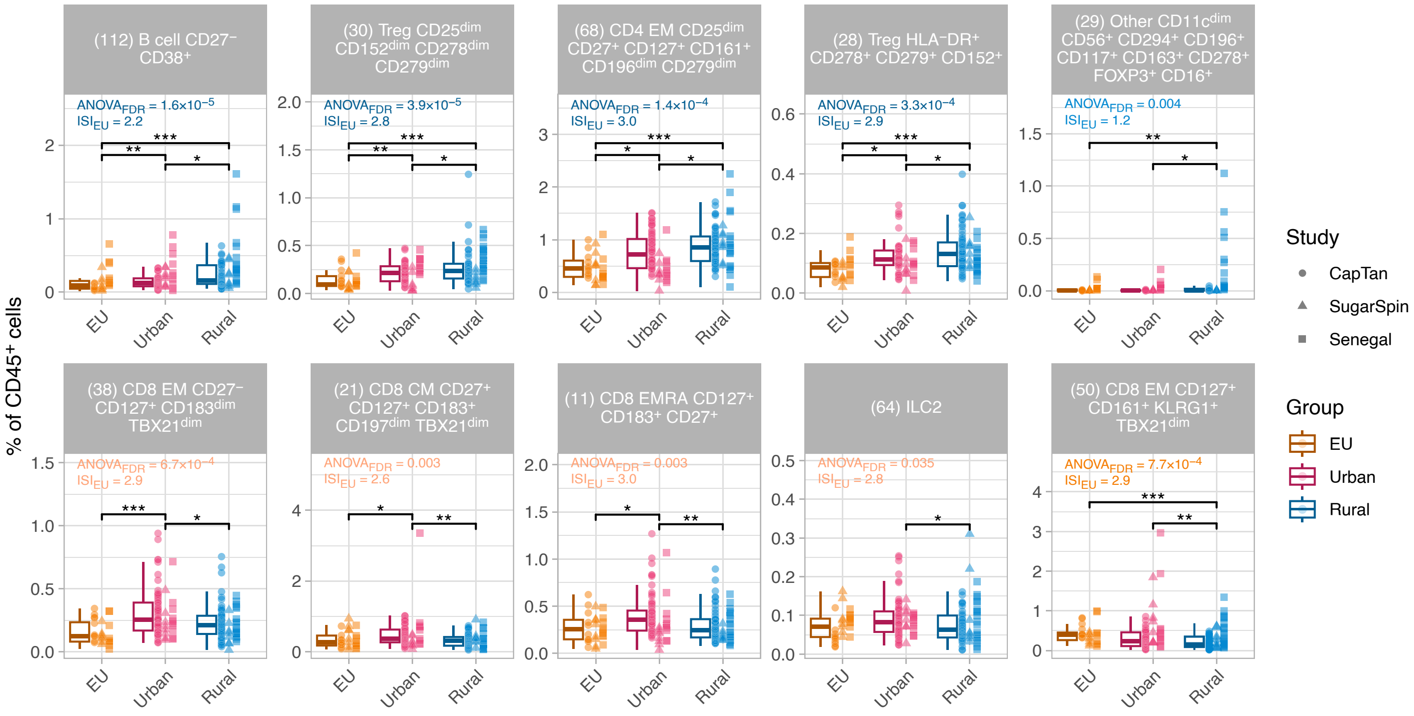

### Supplemental Figure 8

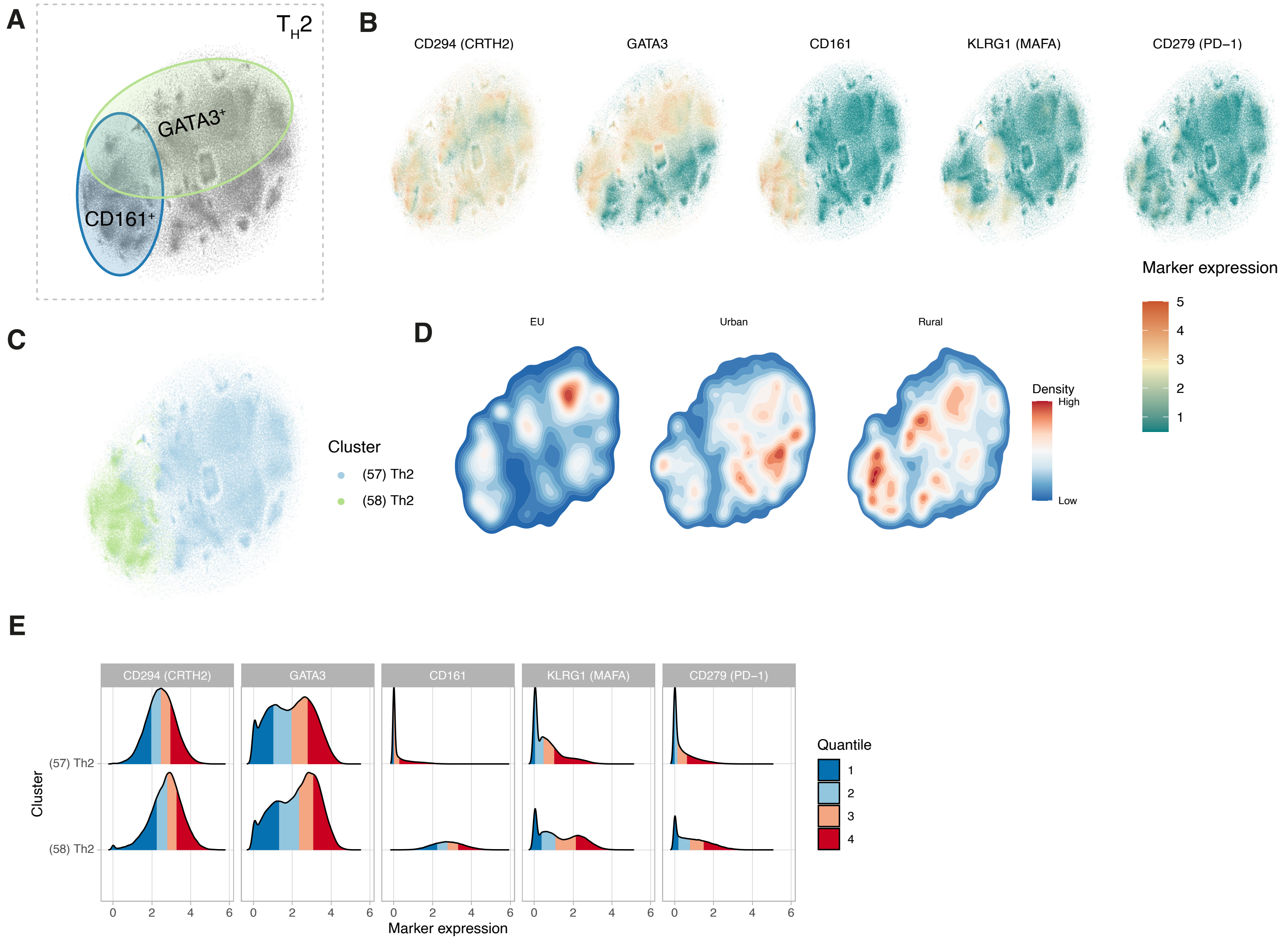

### Supplemental Figure 9

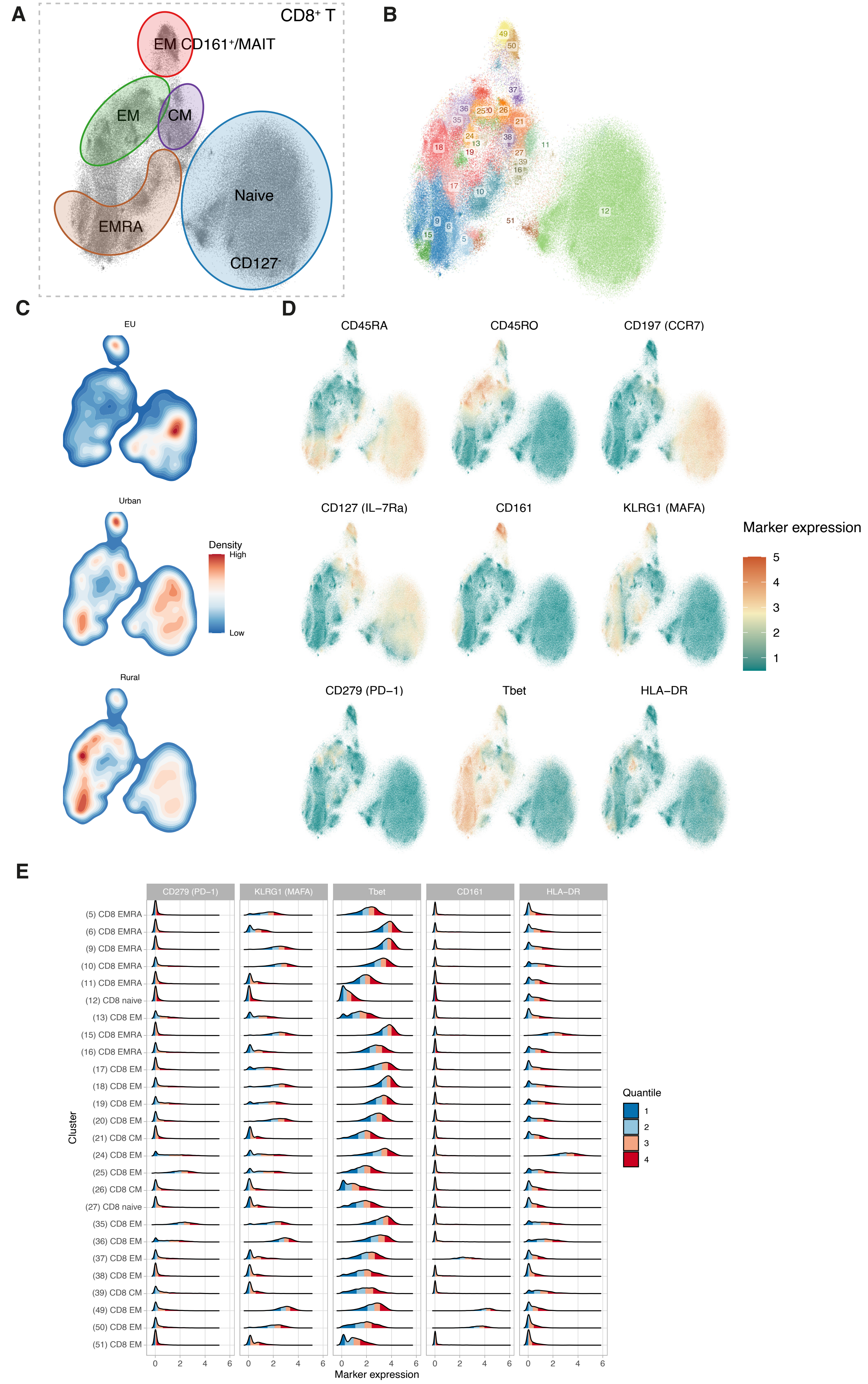

### Supplemental Figure 10

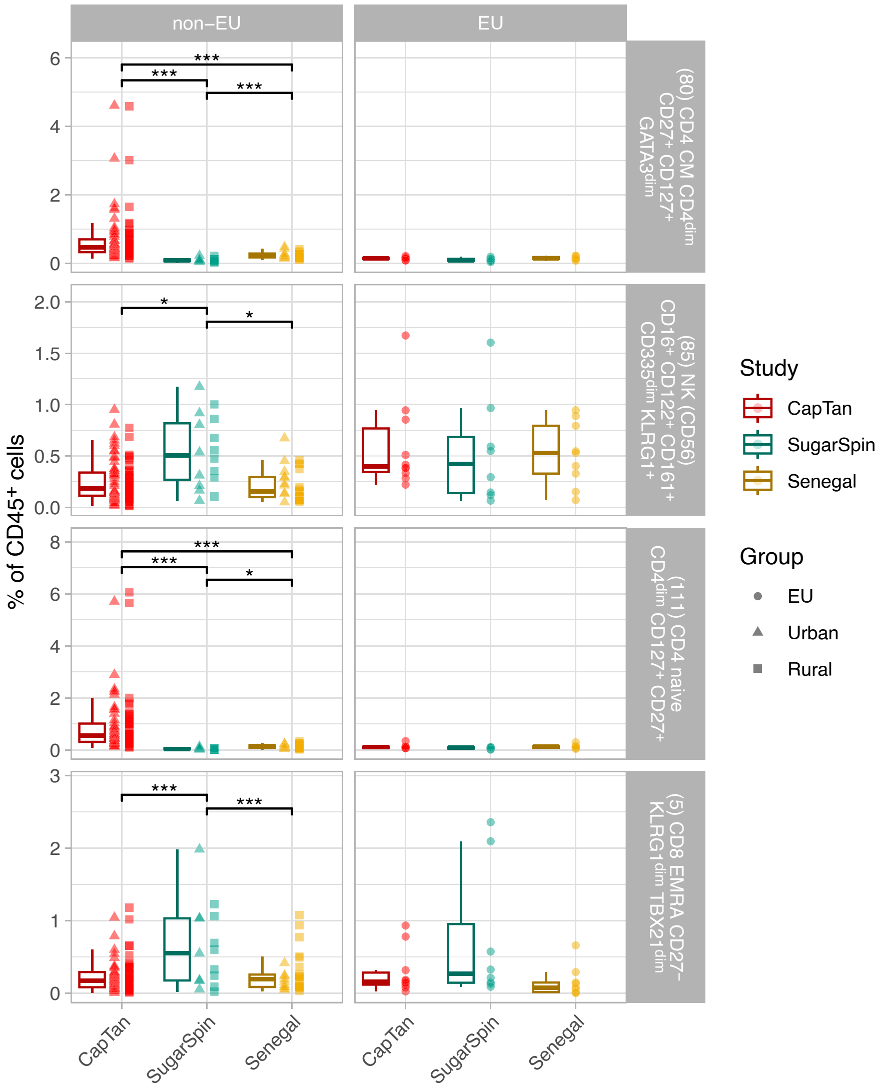
